# Mutant scaling laws reveal that accelerated evolution via gene amplification requires spatially structured population growth

**DOI:** 10.1101/2024.06.24.600326

**Authors:** Natalia L. Komarova, Justin Pritchard, Dominik Wodarz

**Affiliations:** Department of Mathematics, University of California San Diego, La Jolla, CA 92093; Department of Biomedical Engineering, the Pennsylvania State University, State College, PA 16801; Department of Ecology, Behavior & Evolution, University of California San Diego, La Jolla, CA 92093

## Abstract

Principles of evolution in spatially structured expanding populations have recently received much attention, but more work remains to be performed, especially for complex, multi-step evolutionary processes, where mutations are accumulated in an expanding population. A key limitation is that the simulation of spatially explicit stochastic computational models is essential, but not feasible for larger population sizes characteristic of prokaryotic and eukaryotic cell populations. We describe a methodological advance by deriving scaling laws that allow the straightforward prediction of the number of single-hit, double-hit and multi-hit mutants as a function of wild-type population size in spatially expanding populations. While this is a versatile tool to address a range of cutting-edge evolutionary questions, here we apply this methodology to reconcile apparently contradicting data from experimental evolution studies regarding the role of gene amplifications for the emergence of point mutations in bacteria. Applying the scaling laws, we demonstrate that in populations that expand in a 2D or a 3D spatial setting, gene amplifications can significantly promote mutant emergence, and that this is not possible in well-mixed populations. In support of the predictions, experiments that do show accelerated mutant evolution through gene amplifications grew bacteria in spatially restricted lawns, while those that failed to show an effect grew bacteria in non-spatial liquid media.

## 1. Introduction

The accumulation of mutations in expanding cell populations determines the ability of the cells to adapt to changing environments, such as during drug therapy of bacterial infections or tumor cells, or when encountering naturally occurring selection barriers. An extensive theoretical understanding exists about the laws of mutant growth in exponentially expanding populations [1-8], but an equivalent understanding for spatially structured expanding populations remains to be fully worked out. Principles of mutant evolution in spatially structured populations have been elucidated by both computational and experimental approaches [9-24]. Yet, challenges exist on the computational side. Agent-based and similar models, which are typically used to simulate spatial evolutionary processes, become computationally challenging at large population sizes (e.g. when studying bacterial or cancer cell populations), even if more advanced algorithms are used such as the Next Reaction Method [25], Tau-Leaping methods [26] or related approaches [27-29]. For well mixed systems, hybrid simulation methods [30] have been used, in which only relatively small mutant colonies are treated stochastically, while larger sub-populations are simulated deterministically. For spatially explicit simulations, however, the development of such an approach would be highly complex and has not been worked out so far. An alternative way is to develop analytical insights, such as scaling laws, describing how the number of mutants in a growing system scales with the total population based on the system’s dimensionality. A scaling law that connects the number of neutral, single-hit mutants with the colony size was derived previously [9], and the theory was also generalized to neutral and advantageous mutants in 2D and 3D systems [19]. Other types of insightful scaling laws that deal with smooth vs rough fronts, allele surfing, and Muller’s rachet have been derived in [9, 10, 19, 31].

Many biologically relevant evolutionary processes are more complex and involve multiple steps, for which such scaling laws currently do not exist. The aim of this study is to fill this gap. Our work builds on the previous approaches [9, 19] to derive scaling laws for 2- and 3-dimensional population growth for those more complex multi-step evolutionary processes, where intermediate mutants with varying fitness are taken into account. We apply this methodology to predict the number of cells with a target point mutation at population size N, and in particular investigate to what extent an intermediate cell type with an accelerated mutation rate can contribute to the acquisition of the target mutation. Specifically, we consider the role of gene amplifications for mutant evolution. On the one hand, the higher gene copy number can translate into a higher rate of point mutation generation in these genes; on the other hand, the generation and emergence of cells with gene amplifications requires extra rate limiting evolutionary steps. Mathematics can help us resolve this tradeoff and predict the conditions under which gene amplifications enhance the emergence of point mutations in expanding colonies.

Using the growth laws derived here, we find that gene amplifications can only substantially enhance the emergence of target point mutations if population dynamics are characterized by spatial restrictions. This effect is most pronounced for 2D spatially restricted population growth, and is still substantial in 3 dimensions. Interestingly, during growth of well-mixed populations (exponential growth), gene amplifications are not predicted to contribute to the emergence of cells with the target point mutation, unless the fitness advantage of the amplified cells is excessively large and/or the population size reaches biologically unrealistic levels. We use these insights to interpret experimental evolution studies and reconcile apparently contradictory results. We also adapt this methodology do model the role of MMR-deficient somatic cells for mutant generation in expanding populations [32], and find similar results.

## 2. Computer simulations

We consider basic stochastic birth-death processes in expanding populations with mutations. The question under investigation is how many cells that have acquired a specific point mutation (which we will refer to as a “target mutation”) are found in an expanding population at a given size *N*. To address this question, an agent-based model is considered that contains the following populations, see Fig. 1(a,b). (i) <u>Wild-type cells (</u>*<u>Z</u>*_*<u>wt</u>*_), which have not yet acquired any mutation. (ii) <u>Mutant cells (</u>*<u>Z</u>*_*<u>m</u>*_), which have acquired the target mutation due to a replication error of the wild-type population. (iii) <u>Accelerated cells (</u>*<u>Z</u>*_*<u>a</u>*_), which have acquired a gene amplification in the gene under consideration, thus effectively having a higher rate at which the target mutation can be generated. (iv) <u>Mutated accelerated cells (*Z*_*am*_ *<u>+Z</u>*_*<u>ma</u>*_)</u>, which are cells with the higher mutation rate that have acquired the target mutation. Each type of cell is characterized by a rate of cell division and a rate of cell death, denoted by parameters *r* and *d*, respectively, with the appropriate subscript indicating the cell population. Upon division, wild-type cells are assumed to give rise to the target mutation (and hence to cell type *Z*_*m*_) with a rate *μ*. They can also acquire the accelerated phenotype with a rate *p*, giving rise to type *Z*_*a*_ cells. Accelerated cells (*Z*_*a*_) are assumed to produce the target mutation with an elevated rate *μ*_*A*_=*aμ*, giving rise to population *Z*_*am*_. For a cell with gene duplication *a=2*, and larger values of *a* are assumed for higher copy numbers. In addition, mutated cells (*<u>Z</u>*_*<u>m</u>*_) can acquire the accelerated phenotype with a rate *p*, type *<u>Z</u>*<u>_ma_</u>. At every time step, each cell has a chance to divide or die, depending on the assigned probabilities. We compare spatially explicit simulations to a well-mixed system. For spatial simulations, if a division event is chosen, a target spot is selected randomly from all the nearest neighbors (e.g. 8 neighbors for a 2D grid); the offspring is placed there if this target spot is empty, and no division occurs otherwise. If a division occurs, the offspring can be altered with probabilities *μ, μ*_*A*_, or *p*. Non-spatial simulations are performed with Gillespie simulations of the corresponding processes in the context of exponential growth.

**Fig.1.**
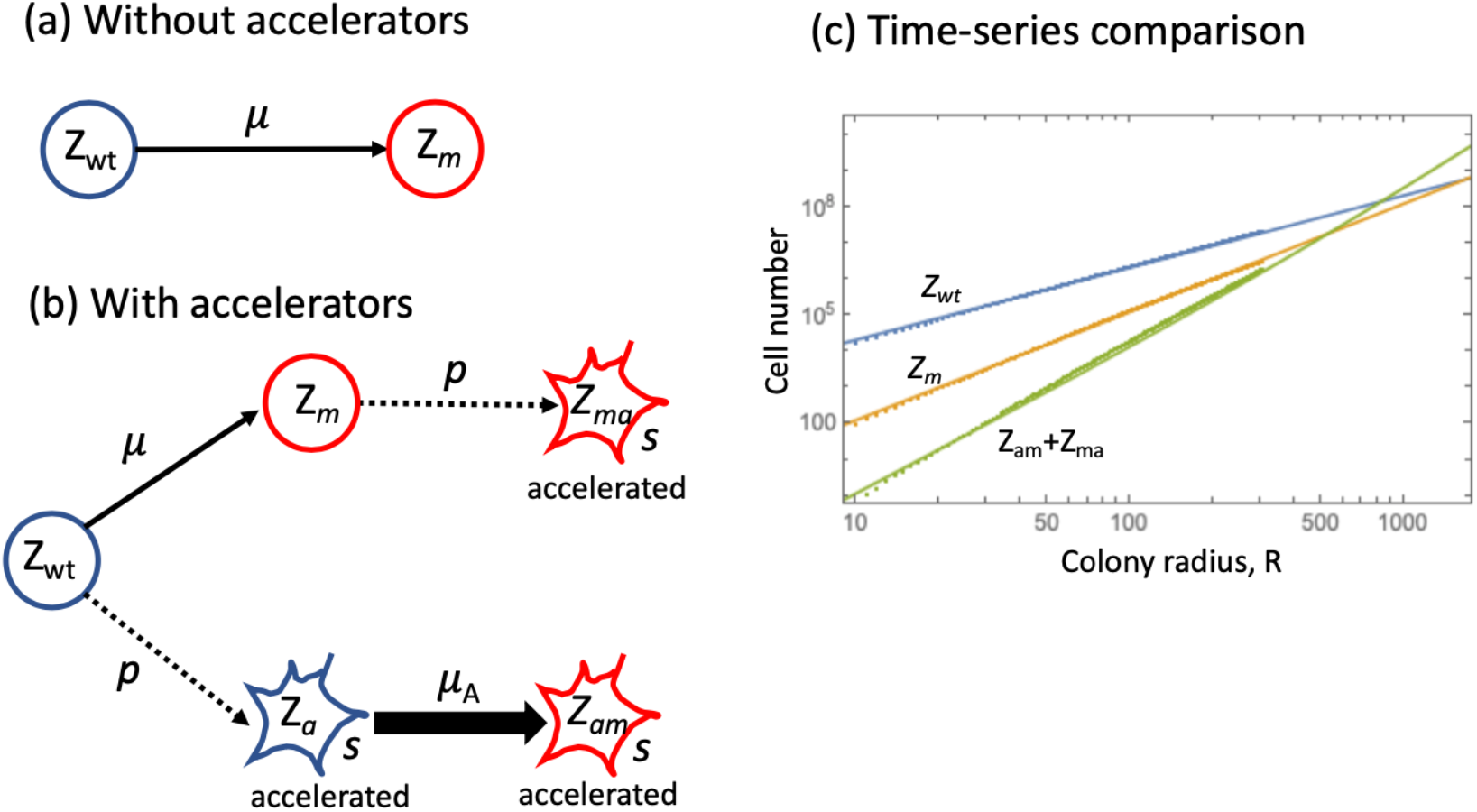
Comparing two pathways to a mutation. (a-b) A schematic showing the pathways leading to the generation of a mutation: (a) in the absence and (b) in the presence of the accelerated phenotype. The accelerated phenotype is denoted by a star shape, and the mutant of interest is colored red. The accelerated mutation rate is denoted by *μ*_*A*_. (c) The number of mutants generated directly (*Z*_*m*_, shown in orange) is plotted together with the number of double-hit mutants generated in the presence of an accelerated type (*Z*_*am*_*+Z*_*ma*_, green) and the number of wild type cells (*Z*_*wt*_, blue) in an expanding 2D colony. Simulations (average of 3,612 independent simulations): dots, theory: lines. Parameters are *s=0*.*01, μ=p =10*^*-4*^, *μ*_*A*_ =2*μ*.

The simulation is started from a single wild-type cell (placed in the center of the grid in the spatial setting). As the simulation proceeds, the population expands according to surface growth (see Fig 2 for the example of a 2D surface growth). During cell divisions, mutated as well as amplified cells are generated with their corresponding rates, and these undergo range expansions that either continue to grow or become enclosed by competing cells over time. In the example shown in Figure 2, both target-mutant (*Z*_*m*_) and amplified cells (*Z*_*a*_) are produced by wild-type cells and undergo range expansions (teal and beige coloring, respectively). Either population can then generate mutated accelerated cells (*Z*_*am*_*+Z*_*ma*_, yellow), which also undergo range expansion dynamics.

**Fig.2.**
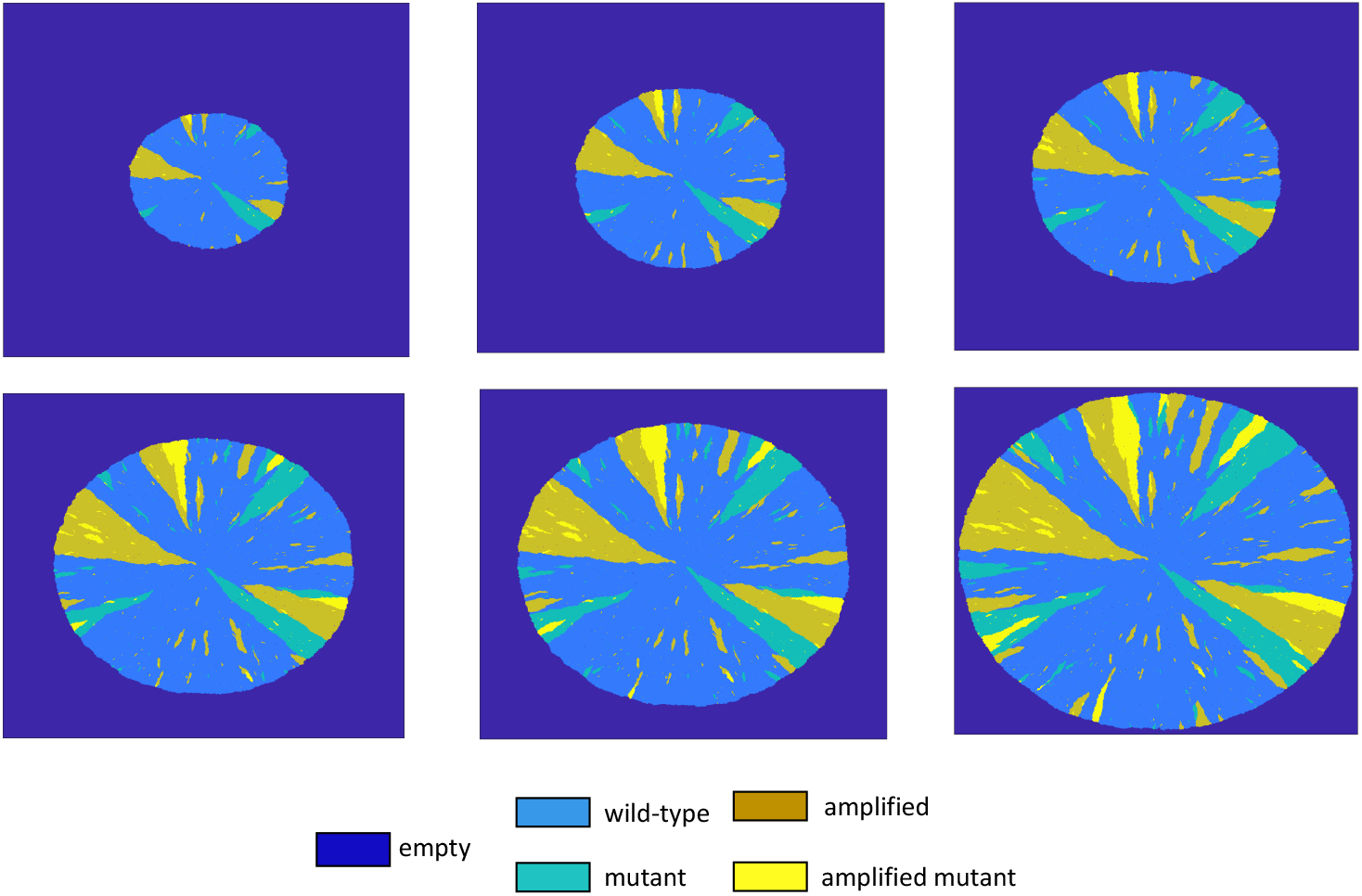
Spatial configuration of a 2D expanding and evolving colony of cells, a typical simulation. The individual graphs represent different snapshots in time, going from left to right top row, followed by left to right bottom row. Color codes are given below the pictures. Parameters are as follows: *r=0*.*985*; *d=0*.*0005*; *s=1*; *p=1*.*5×10*^*-4*^; *μ=1×10*^*-4*^; *a=2*.

To obtain statistics about the number of mutants at a given population size, the simulation is run repeatedly up to a threshold size *N*, and the average number of target mutants at population size *N* is determined (i.e. the sum of populations *Z*_*m*_*+Z*_*am*_*+Z*_*ma*_). Two situations are compared (see the schematic in Fig. 1): simulations in which gene duplications can occur (*p>0*, panel (b)), and simulations where this process is disabled (*p=0*, panel (a)), such that the target mutation can only be generated from the wild-type cell population. For tumor or bacterial cell populations, the number of cells in growing colonies often exceeds what is computationally feasible to simulate in stochastic agent-based models. For illustration purposes, we ran computer simulations for relatively small population sizes and larger mutation rates. An example is seen in Fig 1(c), which assumes that cells with gene duplications have a 1% fitness advantage compared to wild-type cells, and that amplified cells with and without the target mutation have identical fitness. The plot shows the population of wild-type cells (blue) as well as cells that acquire the target mutations directly (*Z*_*m*_, yellow), and the amplified cells that also harbor the target mutation (green). Stochastic simulations results (averaged over a number of independent runs), are shown by dots and show populations up to ∼10^7^ cells. We observe that the single-hit mutants (*Z*_*m*_) seem to increase faster than the wild-type cells, and the double-hit mutants (*Z*_*am*_*+Z*_*ma*_) appear to have an even higher slope in the log-log plot. We next derive mathematical laws of mutant population growth and test them against such simulations. These derived laws are then used to investigate the role of accelerated cells in mutant evolution under biologically realistic population sizes and mutation rates.

## 3. Growth laws of accelerated cells in the spatial model

Computer simulations present evidence that the existence of cells with accelerated mutation rates may contribute significantly to the evolutionary process in spatially distributed systems. They also clearly demonstrate the need for a more general approach, which would allow for a systematic exploration of the evolutionary dynamics under a wide range of parameters. Therefore, we turned to developing scaling laws that govern mutant growth in expanding populations.

### Scaling laws for single mutations

Consider the simplest situation described by the selection-mutation diagram: 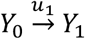, where *Y*_*0*_ denotes the wild-type, *Y*_*1*_ denotes the population of single mutants (with relative fitness *(1+s*_*1*_)), and the mutation rate is given by *u*_*1*_. We will compare two types of spatial growth: a 2D range expansion, where a colony grows in a circular shape, and a 3D range expansion (a spherical colony). We will use variable *R* to characterize the colony size. Denote by R the radius of the colony, where distance is measured in terms of cell diameters. If the total number of cells is denoted by *Y*_*0*_, we have *Y*_0_ ∝ *πR*^2^ in 2D and 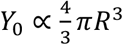 in 3D; in particular, in the absence of cell death, the density of cells is one and the proportionality sign becomes “=“. Assuming so-called surface growth [33], where only cell divisions on the surface contribute to expansion, the mean temporal growth of the population can be described by the following ODEs:

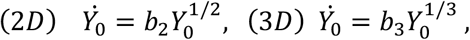

where *b*_*i*_ with 2 ≤ *i* ≤ 3 are constants that include information about the growth rate of cells and also depend on the dimensionality of the system. Consequently, the total number of cells grows according to the following power laws:

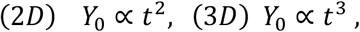

and the linear colony size always grows linearly with time, *R* ∝ *t*.

To include mutant generation, we will assume that mutations are relatively infrequent, such that a single mutation typically gives rise to a clone that does not merge with other clones, and the total mutant number is significantly smaller than the colony size. As a result, it is possible to study mutant clones in isolation. The schematic in figure 3(a) shows a neutral (A) and an advantageous (B) clone growing in a colony undergoing a 2D range expansion. Denote by *n*^*s*^*(r*_*1*_,*r*_*2*_) the expected size of a mutant clone with relative fitness advantage *s*, that was created at colony size *r*_*1*_, and measured at size *r*_*2*_. Further, denote by *M*_*0*_(r) the size of the advancing front, through which the growth occurs: *M*_0_(*r*) ∝ 2*πr* in 2D, and *M*_0_(*r*) ∝ 4*πr*^2^ in 3D. Then the expected number of mutants with fitness advantage *s*_*1*_, in a colony of size *R* can be approximated by: 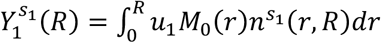, where *u*_*1*_ is the rate at which mutations are generated.

**Fig.3.**
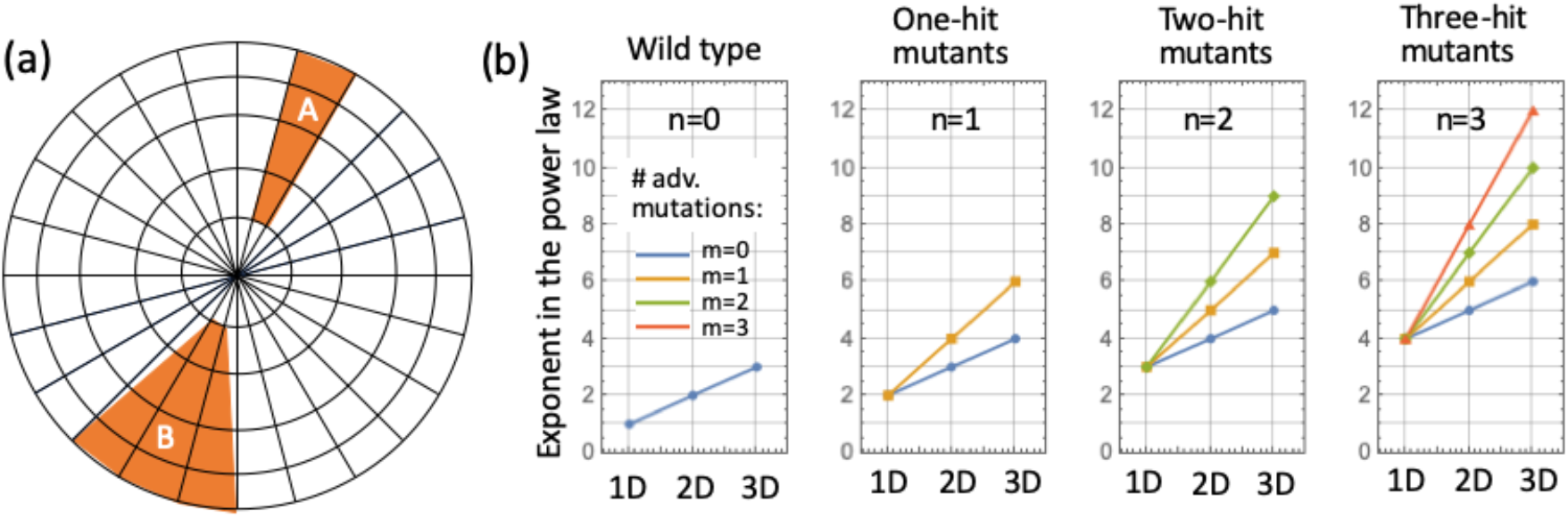
Scaling laws for mutant growth. (a) A schematic showing the spread of a mutant in a 2D range expansion geometry. Mutant A is neutral and its expected domain follows the radial lines. Mutant B is advantageous and its domain boundaries cross radial lines, increasing the arc angle of the mutant region. (b) The scaling laws of eq.(3) and eq.(11) of the Supplement. Each panel shows the exponent in the power law of growth of the population of *n*-hit mutants in 1D, 2D, and 3D geometries. A cell is considered to have an advantage if it has an increase of fitness with respect to the type that generates it (not just the wild type). For the one-, two-, and three-hit mutant scenarios, all the mutants could be neutral (*m*=0, blue line), or one of them could be advantageous (*m*=1, yellow lines). If two (*m*=2) or three (*m*=3) mutations are advantageous, green and red colors are used, respectively. According to the theory, the order in which mutations occur does not matter for the scaling laws.

The expression for the clone size depends on the system dimensionality as well as the mutant fitness relative to the wild type, which is given by *(1+s*_*1*_) with *s*_1_ ≥ 0. It is derived in the supplement for dimensionalities *D=1,2,3*. As a result, we obtain a scaling law for the number of mutants in a colony of linear size *R*, with *s*_1_ ≥ 0 :

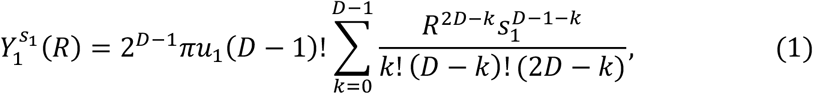

where D=2 or D=3 is the system’s dimensionality. In particular, for neutral mutants (*s*_*1*_=0) the only nonzero term on the right corresponds to *k=D-1*, and we have

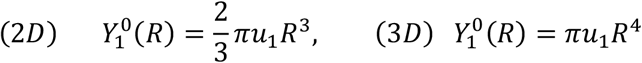

Table 1 shows the power of the leading term for different species in different dimensionalities. For completeness, this table also includes the case D=1, as well as disadvantageous mutants. A more detailed discussion for these laws can be found in [19]; here it suffices to say that each disadvantageous mutant is assumed to give rise to a clone of a constant size, *n*^*s*^*(r*_*1*_,*r*_*2*_)=const, which leads to the results presented in Table 1.

**Table 1.** The leading power of time in the growth laws of different populations in expanding colonies with single mutations, in 1D, 2D, and 3D. See also Fig. 3(b), *m=0*.

| **Single-mutant scaling laws** | 1D | 2D | 3D |
| --- | --- | --- | --- |
| Wild type or disadvantageous mutant ($s_1 < 0$) | $t^1$ | $t^2$ | $t^3$ |
| Neutral mutant ($s_1 = 0$) | $t^2$ | $t^3$ | $t^4$ |
| Advantageous mutant ($s_1 > 0$) | $t^2$ | $t^4$ | $t^6$ |

### Scaling laws for double mutations and beyond

Our goal is to investigate the effect of cells with an increased mutation rate on the evolutionary dynamics. Before such cells can contribute, they have to be created from the wild-type cells, which implies the existence of an additional mutation event. Therefore, we need to develop the growth laws of double-hit mutants. Assume the following selection-mutation diagram: 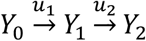, where, as before, *Y*_*0*_ denotes the wild-type, *Y*_*1*_ denotes single mutants (with relative fitness *(1+s*_*1*_)), and *Y*_*2*_ denotes double mutants (with relative fitness *(1+s*_*2*_)). For now we fix the order of events, and assume that type *Y*_*2*_ is generated by a mutation in type *Y*_*1*_. Note that here and below, for each consecutive mutation we measure the resulting fitness with respect to the type that hosts the mutation, and not with respect to the wild type. The rates of mutation are given by *u*_*1*_ and *u*_*2*_ for the first and the second hit, respectively.

The above theory can be extended to include double-hit mutants (see Supplement for details). We assume that double-hit mutants are generated by the single-hit mutants at the edge of expansion. Their expected number is given by: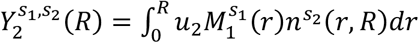, where 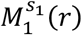 is the number of single mutants of fitness advantage *s*_*1*_ at the edge of expansion for a colony of linear size *r*, and 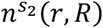 is the size of double-mutant colony of fitness advantage *s*_*2*_ that is generated at size *r* and measured at size *R*. We obtain the following expression for the expected number of double-mutants:

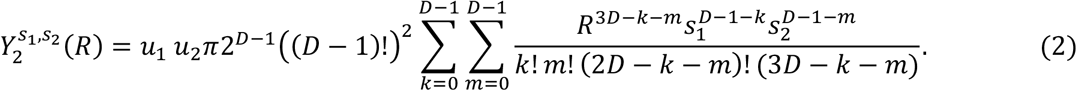

For example, when both mutations are neutral (*s*_*1*_*=s*_*2*_*=0*), the only nonzero contribution comes from the term with *k=m=D-1*, and we obtain

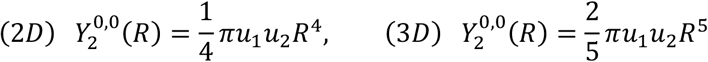

In the Supplement, we provide an explicit formula for any number of consecutive mutations, some or all of which can be advantageous. There are several consequences of the scaling laws that are derived, which we list here.

1. A population of *n*-hit neutral mutants under D dimensional range expansion grows as *R*^*D+n*^, where *R* is the whole colony radius; the power scaling with time is the same, *t*^*D+n*^.
2. Suppose that among *n* sequential hits, *m* hits confer a nonzero advantage with respect to the previous cell type, and the rest of the mutations do not alter the fitness properties of cells that harbor those mutations. Surprisingly, the order in which these advantageous mutations are acquired does not influence the scaling law for the number of *n*-hit mutants, which grow as

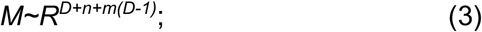

the power scaling with time is the same, *t*^*D+n+m(D-1)*^. The exponents of the scaling law for different values of *n* and *m* are presented in figure 3(b).
3. Suppose that individuals in a population of *n*-hit mutants contain *m* mutations that confer selective advantage to cells (relative to the cell type that acquired the mutation), and denote the corresponding selection coefficient values by *s*_*1*_,*…,s*_*m*_ (where they are listed in a non-decreasing order). If they all have roughly the same order of magnitude (*s*), then the n-hit mutant population behaves as if neutral, until the colony size reaches R∼1/s, at which time it switches to the growth specified in (3). In the case where selective advantage coefficients are different, the growth law increases the power by one at sizes *1/s*_*m*_, *1/s*_*m-1*_,*…, 1/s*_*1*_.

Using this methodology, one can investigate the relative contributions of different pathways to the evolutionary processes in growing populations.

## 4. Using derived growth laws to obtain insights into evolutionary processes in spatially growing populations

Here we show how the analytical expressions for mutant numbers derived above can be used to gain insights into complex evolutionary processes in spatially growing populations. In particular, we ask under what conditions an intermediate population with an increased rate of mutation accumulation can enhance the presence of cells carrying the target point mutation. We first consider gene duplications / amplifications and then also apply this theory to mismatch repair-deficient somatic cells as the intermediate cell population.

### Spatially structured growth is required for gene duplications to significantly accelerate mutant production

To quantify the role of cells with gene duplications for target mutant emergence, we compare the following populations (see Fig. 1): on the one hand, population *Z*_*m*_ in panel (a) is the total number of mutants generated in the absence of mutation acceleration; on the other hand, the sum of the populations *Z*_*am*_*+Z*_*ma*_ represents the number of mutants that are associated with the possibility of accelerating the mutational process. Here, cells *Z*_*am*_ first undergo a mutation that allows for an accelerated mutation rate, and then acquire the target mutation at a higher rate, *μ*_*A*._ Cells *Z*_*ma*_ are created by first acquiring the target mutation and then developing the characteristics of accelerated cells by a second hit. Figure 1(c) compares numerical simulations with the theoretical results for these populations, where for *Z*_*m*_, we used 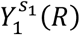 in formula (1) with *u*_*1*_=*μ* and *s*_*1*_*=s;* for *Z*_*ma*_ we used 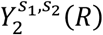 in formula (2) with *u*_*1*_*=μ, u*_*2*_*=p, s*_*1*_*=0, s*_*2*_*=s;* and for *Z*_*am*_ we used 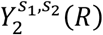 with *u*_*1*_=*p, u*_*2*_*=μ*_*A*_, *s*_*1*_*= s, s*_*2*_*=0*. We can see that the theoretical approximation works well (see further information on validation and applicability in the Supplement).

Using the derived growth laws to predict mutant dynamics for population sizes that are too large to track by computer simulation, we can predict at what colony sizes the cells with duplicated genes make a substantial contribution to target mutant emergence, compared to the number of mutations generated directly by a single point mutation hit from the wild type. Let us define the quantity *Q=(Z*_*am*_*+Z*_*ma*_*)/Z*_*m*_, as the relative contribution of duplicated cells to the total number of mutants created. This quantity has the following straightforward interpretation. Suppose that in the absence of duplicated cells (*p=0*) the number of cells that contains the target mutation in a colony of a given size is *Z*_*m*_. Then in the presence of duplicated cells (*p>0*), the colony of the same size will contain *(1+Q)Z*_*m*_ cells with the target mutation. An alternative measure of the contribution of duplicated cells is 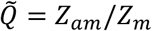, which simply compares the number of cells that acquired the mutation after duplication, with the number of cells that acquired the mutation directly from the wild type. This measure is very similar, and in fact we have 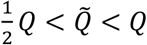, which means that it is somewhere between 50% and 100% of *Q*. If the contribution of duplications in a system is predicted to be significant by measure *Q*, it is also significant by measure 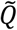. It if is negligible by measure *Q*, it is also negligible by measure 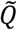.

Assuming that cells with the gene duplication have twice the mutation rate compared to cells without (*μ*_*A*_*=2μ*), Figure 4 compares the behavior of spatial mutant colonies in the cases of 2D and 3D range expansion with a mass-action (exponential growth) system; the methodology for the latter case is standard and is provided in the Supplement. Plotted is the relative contribution of the duplicated cells to the total number of mutants created, *Q=(Z*_*am*_*+Z*_*ma*_*)/Z*_*m*_, for different colony sizes (the horizontal axes). We can see that the relative contribution increases with the colony size. This happens because the net growth rate of the double-mutants is faster; in particular, in the spatial systems it corresponds to an increased exponent in the power law. We also observe that the relative contribution of the duplicated cells is always the largest in 2D (blue), followed by 3D (orange), and both of these are substantially larger than that for the exponentially growing colony (green).

**Fig.4.**
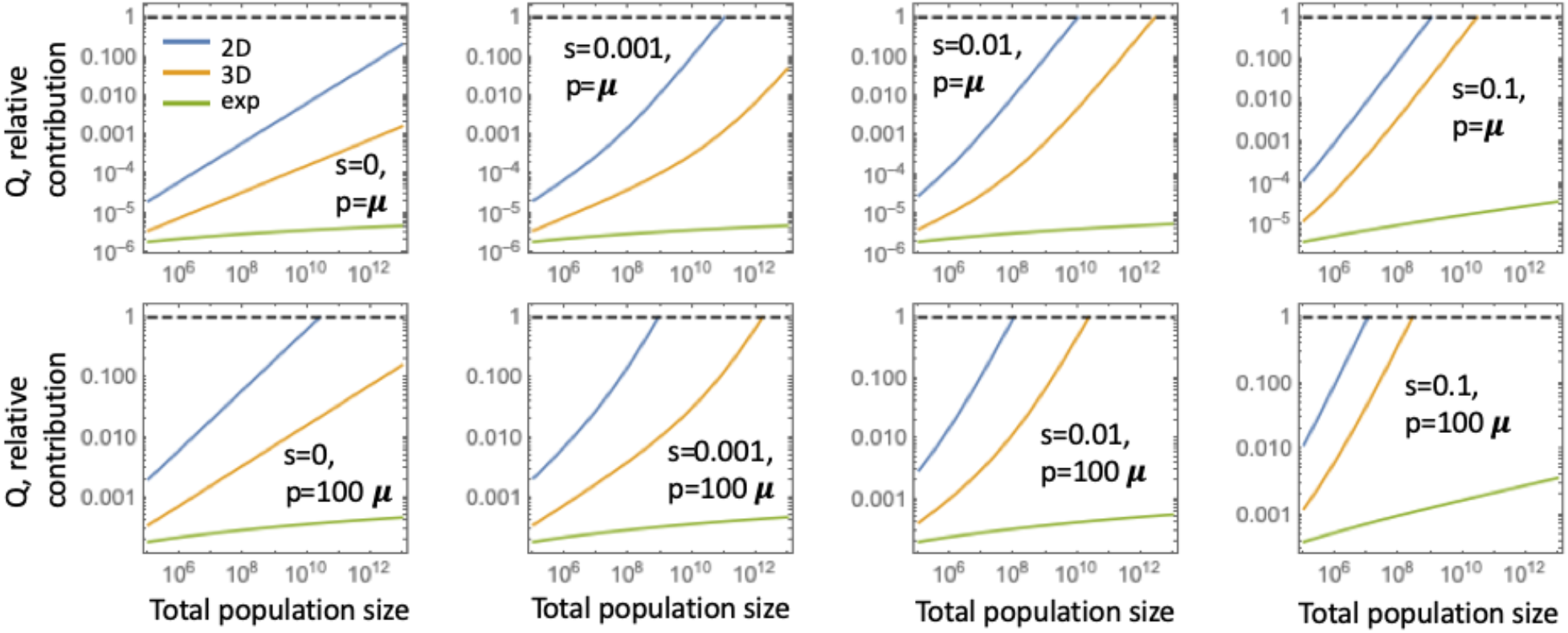
Gene duplication: the relative contribution of the accelerated type to the total number of mutants, *Q=(Z*_*am*_*+Z*_*ma*_*)/Z*_*m*_ (as predicted by the model), is plotted as a function of the total population size. The three lines in each panel correspond to the 2D (blue), 3D (orange), and mass-action (green) systems. We assume that the accelerated mutation rate is *μ*_*A*_*=2μ* (duplications). In the top row, the duplications are generated at rate *p=μ*, and in the bottom row *p=100μ*. The fitness parameter, *s*, increases from left to right as *s=0, s=0*.*001, s=0*.*01*, and *s=0*.*1*. The dashed horizontal line represents *Q=1. μ*_*A*_*=10*^*-7*^.

The striking result is the vast difference in the contribution of the duplicated cells, between the spatial and non-spatial (exponentially growing) systems. In the case of the parameters used in Fig. 4, the increase in the number of target mutants due to gene duplication in the non-spatial model never reaches 1%, even for extremely large populations and assuming that gene duplication happens at a rate that exceeds the base mutation rate a hundredfold and confers a 10% advantage to the cell (see the bottom right panel). In well-mixed populations, gene duplications can only contribute to the accumulation of target point mutations if the fitness of duplicated cells is very large, upwards of 40% (Figure S2). On the other hand, in spatial systems, the increase in the number of target mutants due to duplications can easily reach 100% for much smaller population sizes and much smaller selective advantages of duplicated cells. If gene duplications occur much more often (100-fold in Fig 4, bottom) than point mutations [34], duplicated cells make substantial contribution to target mutant evolution even if the duplicated cells are neutral with respect to the wild-type cells in 2D growth. In the Supplement we present additional graphs where we vary the base mutation rate over a wide range, revealing the same patterns.

### Further amplifications can significantly contribute to mutant production in spatial populations

Beyond duplications, cells can further amplify gene copy number. The schematic of figure 5(a) shows a cascade of amplifications (gene duplications), where each duplication event leads to an increase in the accelerated mutation rate. Here, type *Z*_*1*_ corresponds to type *Z*_*a*_ of figure 1, while types Z_2_, Z_3_, etc represent cells that have experienced 2,3, etc consecutive duplication events. Type *Z*_*1m*_ contains cells that have undergone a single duplication event and also acquired a target mutation (corresponding to the sum *Z*_*am*_*+Z*_*ma*_ in Figure 1(b)), while types *Z*_*2m*_, *Z*_*3m*_, etc are types with multiple duplication events that also harbor the target mutation.

**Fig.5.**
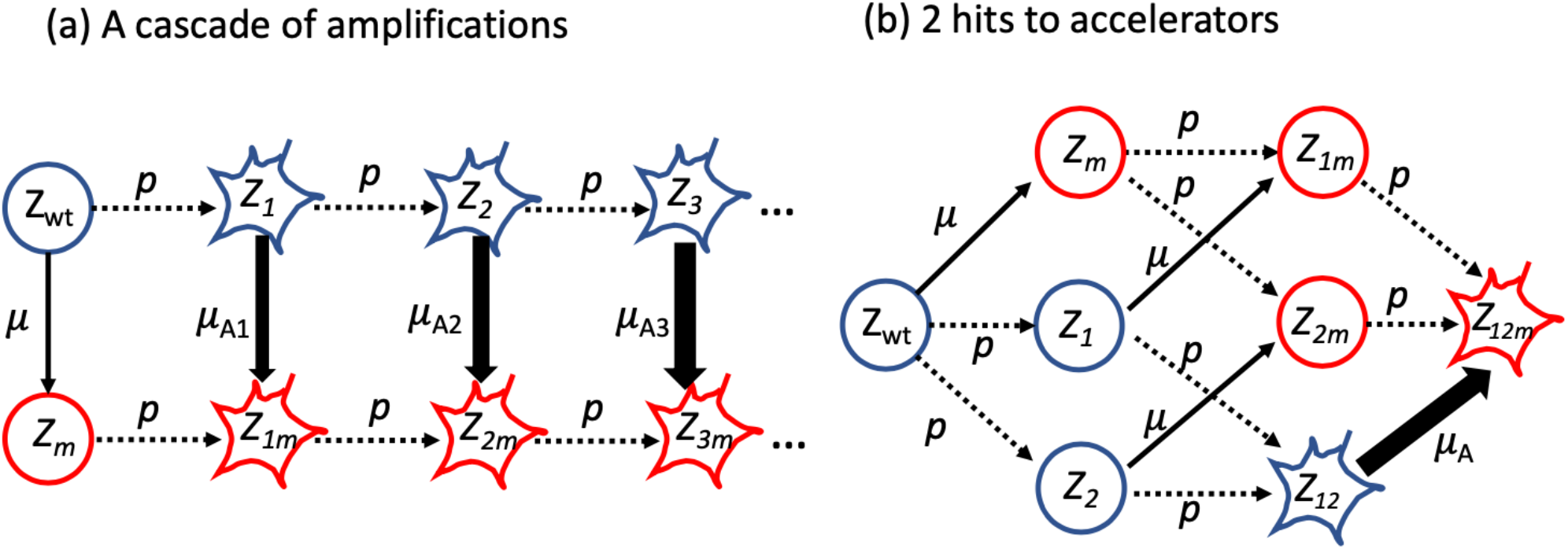
Other selection-mutation networks with mutation accelerations. The accelerated phenotype is denoted by a star shape, and the mutant of interest is colored red. (a) A cascade of amplifications; type *Z*_*k*_ with *k=1,2,…* has experienced *k* gene duplication events in the target gene; type *Z*_*km*_ contains cells with a target mutation that have undergone *k* gene duplication events in the target gene. The increasingly accelerated mutation rates are denoted by *μ*_*A1*_, *μ*_*A2*_, … (b) Two hits to acceleration. Types *Z*_*1*_ and *Z*_*2*_ have the first and the second copy of the MMR gene inactivated, respectively, while the accelerated type *Z*_*12*_ has both copies inactivated. Types *Z*_*1m*_, *Z*_*2m*_, and *Z*_*12m*_ are the same as types *Z*_*1*_ and *Z*_*2*_, and *Z*_*12*_ except they also contain the target mutation.

It is interesting to determine whether or not further duplication events can significantly increase the production of target mutants, beyond the type *Z*_*1m*_. Applying the scaling laws derived here (see Supplement for details) we show that in 2D and 3D systems, further duplication events could make a difference for biologically realistic parameter ranges, if each subsequent event induces additional fitness advantage [34], see figure S7 of the Supplement.

### Spatial population structure is required for the inactivation of an MMR gene in somatic cells (a two-hit event) to contribute significantly to mutant production

MMR deficiency (leading the small-scale genetic instability) is a biologically distinct intermediate cell population that can accelerate the rate of target mutant production in somatic cells, such as colorectal cell types in humans (Fig 5b). In contrast to gene duplications, the generation of the MMR-deficient phenotype in eukaryotic cells requires two mutational events in most cases, because the mismatch repair mechanism needs to be inactivated. In this setting, type *Z*_*12*_ is a cell with both copies of an MMR gene inactivated. It can be produced from the wild type by means of two consecutive inactivation events (through type *Z*_*1*_ or *Z*_*2*_), and is characterized by a significantly (orders of magnitude) increased rate, at which the target mutation can be generated. Hence, although it takes more steps to generate the MMR-deficient phenotype compared to a gene duplication, the gain in the mutation rate is a lot higher. The question is whether for realistic population sizes, this mechanism can contribute to mutant production in expanding populations, such as in colorectal tumors.

Using our scaling laws (see Supplement for details) we can show that, both in 2D and 3D systems, the contribution of MMR-deficient cells can be similar to or larger than direct target mutations. This is in contrast to the non-spatial, exponentially growing systems, where the effect of these longer pathways is negligible, see Fig. S5 of the Supplement.

## 5. Discussion

Stochastic computer simulations that involve evolutionary processes are computationally very “expensive,” and are difficult to perform under realistic population sizes. In the case of spatially expanding populations that produce neutral or advantageous mutants, we developed an approach that is based on newly derived scaling laws for these populations. Using these laws, we can study multi-step evolutionary processes that involve neutral and/or advantageous mutations, and quantify the role played by various processes in the evolutionary transformation experienced by populations. In particular, this methodology allows reasoning about the relative likelihood of different pathways and suggest which events may be causal, given the patterns that are observed. This can be done under realistic assumptions about the mutation rates and population sizes, which is difficult to do using traditional methods of stochastic agent-based modeling.

The scaling laws derived here predict the expected size of a mutant population created by an expanding colony of cells/organisms, both in 2D and 3D range expansion. Results include one-hit mutants, as well as mutants that are created by a longer chain of mutation events, which may include both neutral and advantageous hits. These theoretical results were applied to study the role of gene duplications / amplifications for target mutant emergence, and further expanded to describe the contribution of MMR-deficient somatic cells to mutant production. An emerging theme was that spatial population structure might be crucial for intermediate cell genotypes with elevated mutation rates to contribute substantially to target mutant generation. The effect was strongest for 2D spatial population structures, still substantial for 3D spatial population structures, but largely negligible for well-mixed populations.

Our work adds to an extensive and expanding literature on evolutionary dynamics in spatially structured populations. In recent years, spatial evolutionary dynamics have garnered significant mathematical interest, both computationally and theoretically. Agent-based models have been used to show that the number of mutants at a given threshold size in spatially growing populations are larger than at the same size in a well-mixed system, due to the emergence of jackpot mutations [9]. Further work has shown that this pattern can be more complex for disadvantageous mutants and depend on the parameter in which the disadvantage is expressed [19]. These studies also defined mutant scaling laws for single-hit mutants, which formed the basis for our work., see also [10-12]. Other theoretical work [35] examined a spatially distributed population where mutations arose and spread to neighboring locations, calculating the time distribution required for an individual to acquire a certain number of mutations. Deterministic equations for stochastic spatial games were developed under the assumption of long-range interactions [36]. Another study compared a death-birth process on a lattice with the dynamics of the replicator equation, revealing that spatial factors significantly altered the evolutionary outcomes [37]. These studies build on a rich tradition of mathematical research into spatial evolutionary stochastic processes, utilizing both analytical approaches and partial differential equation (PDE) modeling to study trait evolution in spatial contexts [38-42].

Spatial evolutionary processes have also been investigated specifically in the context of cancer, with implications for tumor progression and expansion [22], for spatial heterogeneity and tumor sampling [43], and for the evolution and management of drug resistance [15, 44].

### Modeling results help interpret experimental evolution data on the role of copy number variation

Gene amplifications are known to be important drivers of evolutionary processes [34, 45]. While amplifications by themselves can be costly to cells, they can help to increase fitness under stressful conditions. An extensively studied and debated system are *Escherichia coli* cells with a mutation in the lac operon (*lac*^*-*^) [46-49], which renders them almost unable to grow on lactose as the carbon and energy source (only about 2% enzyme activity compared to wild-type). When a large bacterial population is plated on lactose, populations of revertant cells (*lac*^*+*^) emerge. While there has been debate about the mechanisms underlying this evolution, a picture has emerged, in which gene amplifications play a major role. It is thought that the originally plated cells contain small subpopulations with a lac operon duplication. This allows the cells to grow slowly on lactose, and subsequent further gene amplifications allow for more efficient growth. At the same time, the increased gene copy number results in a higher chance to generate a revertant mutant that allows the bacteria to grow on lactose even without the gene amplification. Since gene amplifications by themselves are costly to cells, the increased copy numbers are subsequently lost. This represents the type of evolutionary process that we studied mathematically here (except that we did not explicitly take into account loss of amplifications, and the target mutation could be any mutation in the amplified sequence, not just a mutation that functionally replaces an amplification). The same kind of patterns have also been documented for Salmonella enterica [50]. In all these experiments, the bacteria grew as lawns, which represent 2D spatial growth dynamics. According to our model, this is the population structure in which it is most likely to detect cells with point mutations that were generated in cells with gene amplifications. Similar evolutionary processes are thought to occur under other conditions of stress, including antibiotic resistance evolution, heat stress, or salt stress [51, 52].

The role of gene duplications and amplifications for the rate of mutant evolution in *E. coli* has also been investigated in experiments in which bacteria grew in liquid medium [53]. This represents a more mixed and less spatial system. A synthetic genetic reporter system in *E. coli* was used to distinguish between copy number and point mutations, and focused on the early phase of the evolutionary dynamics. The system involved an intact endogenous *galK* gene with a random promoter system. When *E. coli* is grown in the presence of galactose, there is selection pressure for increased *galK* expression, which can occur either through gene amplification or through point mutation. Among the experimental conditions investigated in this study, the low galactose environment [53] is the most applicable to our model, because the point mutations generated in amplified cells did not increase the bacterial fitness beyond the advantage already attained through amplification. In this system it was found that bacteria carried either one or the other type of alteration, but not both together. The explanation given for this observation was based on clonal interference between adaptive amplifications and point mutations [53]. It is interesting to point out, however, that our model also predicts that lack of co-occurrence of gene amplifications and point mutations in this system because the bacteria were grown in liquid medium. In a mixed system, our model suggests that to observe the co-occurrence of point mutations and gene amplifications in cells, the population size would have to grow to unrealistically large sizes, or the fitness advantage of the amplified cells would have to be extremely large. Hence, it is possible that the lack of point mutations arising in amplified cells in these experiments is the result of the basic growth laws of amplified cells in a mixed system (as opposed to a spatial system), without the need to invoke further complexities. It would be interesting to repeat this study in a setting where bacterial growth is more spatially restricted.

The result that spatial population structure greatly promotes the ability of gene amplifications to contribute to the generation of point mutations is important for a detailed interpretation and comparison of experimental results across different systems. It also has important implications for cancer. Since the growth of tumors (especially solid tumors) is characterized by strong spatial restrictions, gene amplifications likely accelerate the accumulation of point mutations in the amplified genes, which can in turn drive the adaptation of tumor cells to overcome natural selective barriers or to become resistant to therapies. While gene amplifications readily occur in cancer genomes [54], this might be especially important for extrachromosomal DNA [55, 56] (ecDNA). Oncogene amplifications occur on ecDNA, and this can promote drug resistance and disease progression through further evolutionary processes, including the acquisition of point mutations in amplified genes.

Apart from gene amplifications, we also considered MMR-deficient cells as an intermediate state with an elevated mutation rate. While MMR-deficiency occurs in bacteria, a more relevant system in the context of our model is MMR-deficiency in somatic cells, such as in colorectal cancer cells. This has been shown to accelerate colorectal cancer evolution through microsatellite instability. The MMR-deficient phenotype in this system typically requires the inactivation of the MMR gene, which involves two separate mutational events. While this slows down the evolutionary pathway via MMR-deficiency, our model predicts that in the context of spatially structured cell growth, the MMR-deficient cells that are generated in a growing population can still substantially contribute to the generation of target mutations, but only if the MMR-deficient cells have a selective advantage. This is likely to be true for colorectal cancer.

Interestingly, the gene inactivation leading to MMR-deficiency also leads to a reduction in the rate at which cells undergo apoptotic cell death, which provides these cells with an immediate selective advantage that can outweigh fitness reduction through the accelerated generation of deleterious mutants [32]. According to our model, this intrinsic fitness advantage of MMR-deficient cells, coupled with the spatial growth of colorectal tumors, allows MMR-deficient cells to drive tumor evolution (and this would not occur if these conditions were violated).

## 6. Conclusions

Through the derivation of mutant scaling laws in spatially structured and expanding populations, we have provided a tool to study various multi-step evolutionary processes during spatial population growth without the restrictions posed by computationally intensive agent-based models. Hence, we can answer evolutionary questions in the context of large populations and low mutation rates, which is relevant for bacterial and cancer cell populations. We have applied this methodology to study the role of intermediate cell types with accelerated mutation rates for the acquisition of target point mutations, particularly focusing on the role of gene duplications / amplifications, and further expanding this to MMR-deficiency in eukaryotic cells. This methodology, however, is more broadly applicable, and can be used to answer questions concerned with the evolution of complex traits in growing populations. This has not only implications for evolutionary biology, but also for biomedical questions such as the evolution of therapy resistance, immune escape, and aspects of disease development.

## References

1. Luria SE, Delbruck M. Mutations of bacteria from virus sensitivity to virus resistance. Genetics. 1943;28:491–511.

2. Zheng Q. Progress of a half century in the study of the Luria-Delbruck distribution. Mathematical biosciences. 1999;162(1-2):1-32. PubMed PMID: 10616278.

3. Dewanji A, Luebeck EG, Moolgavkar SH. A generalized Luria-Delbruck model. Mathematical biosciences. 2005;197(2):140-52. PubMed PMID: 16137718.

4. Komarova NL, Wu L, Baldi P. The fixed-size Luria–Delbruck model with a nonzero death rate. Mathematical biosciences. 2007;210(1):253–90.

5. Kepler TB, Oprea M. Improved inference of mutation rates: I. An integral representation for the Luria–Delbrück distribution. Theoretical population biology. 2001;59(1):41–8.

6. Goldie JH, Coldman AJ. A model for resistance of tumor cells to cancer chemotherapeutic agents. Mathematical biosciences. 1983;65:291–307.

7. Goldie JH, Coldman AJ. Drug resistance in cancer: mechanisms and models: Cambridge University Press; 1998.

8. Komarova NL, Wodarz D. Drug resistance in cancer: principles of emergence and prevention. Proceedings of the National Academy of Sciences of the United States of America. 2005;102(27):9714–9. Epub 2005/06/28. doi: 0501870102 [pii] 10.1073/pnas.0501870102. PubMed PMID: 15980154; PubMed Central PMCID: PMC1172248.

9. Fusco D, Gralka M, Kayser J, Anderson A, Hallatschek O. Excess of mutational jackpot events in expanding populations revealed by spatial Luria-Delbruck experiments. Nature communications. 2016;7:12760. doi: 10.1038/ncomms12760. PubMed PMID: 27694797; PubMed Central PMCID: PMC5059437.

10. Gralka M, Hallatschek O. Environmental heterogeneity can tip the population genetics of range expansions. Elife. 2019;8. Epub 2019/04/13. doi: 10.7554/eLife.44359. PubMed PMID: 30977724; PubMed Central PMCID: PMCPMC6513619.

11. Gralka M, Stiewe F, Farrell F, Mobius W, Waclaw B, Hallatschek O. Allele surfing promotes microbial adaptation from standing variation. Ecology letters. 2016;19(8):889–98. Epub 2016/06/17. doi: 10.1111/ele.12625. PubMed PMID: 27307400; PubMed Central PMCID: PMCPMC4942372.

12. Otwinowski J, Krug J. Clonal interference and Muller’s ratchet in spatial habitats. Physical biology. 2014;11(5):056003. Epub 2014/08/27. doi: 10.1088/1478-3975/11/5/056003. PubMed PMID: 25156977.

13. Lavrentovich MO, Wahl ME, Nelson DR, Murray AW. Spatially Constrained Growth Enhances Conversional Meltdown. Biophysical journal. 2016;110(12):2800–8. Epub 2016/06/23. doi: 10.1016/j.bpj.2016.05.024. PubMed PMID: 27332138; PubMed Central PMCID: PMCPMC4919427.

14. Waclaw B, Bozic I, Pittman ME, Hruban RH, Vogelstein B, Nowak MA. A spatial model predicts that dispersal and cell turnover limit intratumour heterogeneity. Nature. 2015;525(7568):261–4. doi: 10.1038/nature14971. PubMed PMID: 26308893; PubMed Central PMCID: PMC4782800.

15. Aif S, Appold N, Kampman L, Hallatschek O, Kayser J. Evolutionary rescue of resistant mutants is governed by a balance between radial expansion and selection in compact populations. Nature communications. 2022;13(1):7916.

16. Yu Q, Gralka M, Duvernoy M-C, Sousa M, Harpak A, Hallatschek O. Mutability of demographic noise in microbial range expansions. The ISME journal. 2021;15(9):2643–54.

17. Paulose J, Hallatschek O. The impact of long-range dispersal on gene surfing. Proceedings of the National Academy of Sciences. 2020;117(14):7584–93.

18. Birzu G, Matin S, Hallatschek O, Korolev KS. Genetic drift in range expansions is very sensitive to density dependence in dispersal and growth. Ecology letters. 2019;22(11):1817–27.

19. Wodarz D, Komarova NL. Mutant Evolution in Spatially Structured and Fragmented Expanding Populations. Genetics. 2020;216(1):191–203.

20. Schreck CF, Fusco D, Karita Y, Martis S, Kayser J, Duvernoy M-C, et al. Impact of crowding on the diversity of expanding populations. Proceedings of the National Academy of Sciences. 2023;120(11):e2208361120.

21. Kayser J, Schreck CF, Gralka M, Fusco D, Hallatschek O. Collective motion conceals fitness differences in crowded cellular populations. Nature ecology & evolution. 2019;3(1):125–34.

22. Noble R, Burri D, Le Sueur C, Lemant J, Viossat Y, Kather JN, et al. Spatial structure governs the mode of tumour evolution. Nature ecology & evolution. 2022;6(2):207–17.

23. Colyer B, Bak M, Basanta D, Noble R. A seven-step guide to spatial, agent-based modelling of tumour evolution. Evolutionary applications. 2024;17(5):e13687.

24. West J, Schenck RO, Gatenbee C, Robertson-Tessi M, Anderson AR. Normal tissue architecture determines the evolutionary course of cancer. Nature communications. 2021;12(1):2060.

25. Gibson MA, Bruck J. Efficient exact stochastic simulation of chemical systems with many species and many channels. The journal of physical chemistry A. 2000;104(9):1876–89.

26. Cao Y, Gillespie DT, Petzold LR. Efficient step size selection for the tau-leaping simulation method. Journal of Chemical Physics. 2006;124(4). doi: Artn 04410910.1063/1.2159468. PubMed PMID: WOS:000234979300010.

27. Gillespie DT. Stochastic simulation of chemical kinetics. Annual Review of Physical Chemistry. 2007;58:35–55. doi: 10.1146/annurev.physchem.58.032806.104637. PubMed PMID: WOS:000246652300003.

28. Salis H, Kaznessis Y. Accurate hybrid stochastic simulation of a system of coupled chemical or biochemical reactions. Journal of Chemical Physics. 2005;122(5). doi: Artn 054103 10.1069/1.1835951. PubMed PMID: WOS:000226880100004.

29. Wilkinson DJ. Stochastic modelling for systems biology: CRC press; 2011.

30. Rodriguez-Brenes IA, Komarova NL, Wodarz D. The role of telomere shortening in carcinogenesis: A hybrid stochastic-deterministic approach. Journal of theoretical biology. 2019;460:144–52. doi: 10.1016/j.jtbi.2018.09.003. PubMed PMID: 30315815; PubMed Central PMCID: PMC6234035.

31. Granelli-Piperno A, Shimeliovich I, Pack M, Trumpfheller C, Steinman RM. HIV-1 selectively infects a subset of nonmaturing BDCA1-positive dendritic cells in human blood. Journal of immunology. 2006;176(2):991-8. PubMed PMID: 16393985.

32. Madden-Hennessey K, Gupta D, Radecki AA, Guild C, Rath A, Heinen CD. Loss of mismatch repair promotes a direct selective advantage in human stem cells. Stem cell reports. 2022;17(12):2661–73.

33. Rodriguez-Brenes IA, Komarova NL, Wodarz D. Tumor growth dynamics: insights into evolutionary processes. Trends in ecology & evolution. 2013;28(10):597–604. doi: 10.1016/j.tree.2013.05.020. PubMed PMID: 23816268.

34. Andersson DI, Hughes D. Gene amplification and adaptive evolution in bacteria. Annual review of genetics. 2009;43:167–95.

35. Foo J, Leder K, Schweinsberg J. Mutation timing in a spatial model of evolution. Stochastic Processes and their Applications. 2020;130(10):6388–413.

36. Hwang SH, Katsoulakis M, Rey-Bellet L. Deterministic equations for stochastic spatial evolutionary games. Theoretical Economics. 2013;8(3):829–74.

37. Evilsizor S, Lanchier N. Evolutionary games on the lattice: death-birth updating process. 2016.

38. Champagnat N, Méléard S. Invasion and adaptive evolution for individual-based spatially structured populations. J Math Biol. 2007;55:147–88.

39. Roca CP, Cuesta JA, Sánchez A. Effect of spatial structure on the evolution of cooperation. Physical Review E. 2009;80(4):046106.

40. Ovaskainen O, Finkelshtein D, Kutoviy O, Cornell S, Bolker B, Kondratiev Y. A general mathematical framework for the analysis of spatiotemporal point processes. Theoretical ecology. 2014;7:101–13.

41. Ibsen-Jensen R, Chatterjee K, Nowak MA. Computational complexity of ecological and evolutionary spatial dynamics. Proceedings of the National Academy of Sciences. 2015;112(51):15636–41.

42. Chen Y-T. Wright–Fisher diffusions in stochastic spatial evolutionary games with death– birth updating. 2018.

43. Opasic L, Zhou D, Werner B, Dingli D, Traulsen A. How many samples are needed to infer truly clonal mutations from heterogenous tumours? BMC cancer. 2019;19:1–11.

44. Strobl MA, Gallaher J, West J, Robertson-Tessi M, Maini PK, Anderson AR. Spatial structure impacts adaptive therapy by shaping intra-tumoral competition. Communications medicine. 2022;2(1):46.

45. Elliott KT, Cuff LE, Neidle EL. Copy number change: evolving views on gene amplification. Future microbiology. 2013;8(7):887–99.

46. Hastings P, Bull HJ, Klump JR, Rosenberg SM. Adaptive amplification: an inducible chromosomal instability mechanism. Cell. 2000;103(5):723–31.

47. Hersh MN, Ponder RG, Hastings P, Rosenberg SM. Adaptive mutation and amplification in Escherichia coli: two pathways of genome adaptation under stress. Research in Microbiology. 2004;155(5):352–9.

48. Hendrickson H, Slechta ES, Bergthorsson U, Andersson DI, Roth JR. Amplification– mutagenesis: Evidence that “directed” adaptive mutation and general hypermutability result from growth with a selected gene amplification. Proceedings of the National Academy of Sciences. 2002;99(4):2164–9.

49. Kugelberg E, Kofoid E, Reams AB, Andersson DI, Roth JR. Multiple pathways of selected gene amplification during adaptive mutation. Proceedings of the National Academy of Sciences. 2006;103(46):17319–24.

50. Slechta ES, Bunny KL, Kugelberg E, Kofoid E, Andersson DI, Roth JR. Adaptive mutation: general mutagenesis is not a programmed response to stress but results from rare coamplification of dinB with lac. Proceedings of the National Academy of Sciences. 2003;100(22):12847–52.

51. Yona AH, Manor YS, Herbst RH, Romano GH, Mitchell A, Kupiec M, et al. Chromosomal duplication is a transient evolutionary solution to stress. Proceedings of the National Academy of Sciences. 2012;109(51):21010–5.

52. Kondrashov FA. Gene duplication as a mechanism of genomic adaptation to a changing environment. Proceedings of the Royal Society B: Biological Sciences. 2012;279(1749):5048–57.

53. Tomanek I, Guet CC. Adaptation dynamics between copy-number and point mutations. Elife. 2022;11:e82240.

54. Albertson DG. Gene amplification in cancer. TRENDS in Genetics. 2006;22(8):447–55.

55. Lange JT, Rose JC, Chen CY, Pichugin Y, Xie L, Tang J, et al. The evolutionary dynamics of extrachromosomal DNA in human cancers. Nature genetics. 2022;54(10):1527–33.

56. Wu S, Bafna V, Chang HY, Mischel PS. Extrachromosomal DNA: an emerging hallmark in human cancer. Annual Review of Pathology: Mechanisms of Disease. 2022;17:367–86.

