## Supplemental information for "Mutant scaling laws reveal that accelerated evolution via gene amplification requires spatially structured population growth"

**4 An example of a model of fitness in a 2D flat expansion 17**

### 1 Scaling laws for mutant generation and growth in expanding populations

#### 1.1 Model assumptions and fitness

In this model, we assume that the population of wild-type individuals is expanding according to a simple deterministic law. For 1D, the colony has the shape of a 1D interval of length $R$; in 2D it is a circle of radius $R$ and in 3D it is a sphere of radius $R$. The growth takes place by adding consecutive layers of cells; these layers will be identified by their coordinate $r$.

We assume the existence of a mutation sequence, where mutations are acquired by a cell in a given order, with mutation rates $u_1$, $u_2$, etc. We further assume that a cell type that contains the first $i$ mutations is characterized by a fitness advantage (relative to the type that generated the mutation, not the wild type) of $s_i$.

Let us denote by $M_0(r)$ the area (in the subspace perpendicular to the direction of expansion) of the wild-type at size $r$,

$$M_0(r) = C_D(2r)^{D-1},$$

where $C_D$ is a constant that depends on the system's dimensionality:

$$C_D = \begin{cases} 1, & 1D, \\ \pi, & 2D, \\ \pi, & 3D. \end{cases}$$

The total number of wild-type cells at size $R$ is given by

$$Y_0(R) = \int_0^R M_0(r)\,dr = \frac{2^{D-1} C_D R^D}{D}. \tag{1}$$

We assume that mutant clones originate at the front of the expanding parent population and advance forward with it. Let us define as $a^{s_1}(r_1, r_2)$ the front area of a mutant colony of advantage $s_1$, which originated at size $r_1$ and is measured at size $r_2$:

$$a^{s_1}(r_1, r_2) = \left(\frac{r_2}{r_1}\right)^{D-1} (1 + s_1(r_2 - r_1))^{D-1}. \tag{2}$$

To justify this expression, we consider several cases.

- *A 2D flat front expansion.* This is the simplest geometry, which was considered e.g. in [1], but it is not explicitly included in the main text of this study. We use it to motivate and transition to other geometries. In this system, wild-type cells in 2D grow as a planar front in one direction, creating a rectangle of a constant width $W \gg 1$ and a changing length, $R$, with $R \propto t$. Single-hit mutants are created at rate $Wu_1$ per layer. If a neutral mutant is created at layer $r_1$, it creates a clone of width 1. If however the mutant cells have a selective advantage, each consecutive layer will contain $s_1$ more cells, see the discussion in Section 4. As a consequence, the width of the mutant clone created at size $r_1$ and measured at size $r_2$ is given by

$$1 + s_1(r_2 - r_1). \tag{3}$$

- *A 2D range expansion.* The colony expands outward layer by layer, and at the layer of radius $r$, the probability to create a mutant is $2\pi r_1 u_1$. A neutral mutant clone created at radius $r_1$ will on average expand outwards in such a way that the fraction of mutant cells in each layer is constant and equal to $\frac{1}{2\pi r_1}$, the initial fraction. The length of a neutral mutant front at colony radius $r_2$ is given by $\frac{2\pi r_2}{2\pi r_1} = \frac{r_2}{r_1}$. For advantageous mutants, the fraction of mutants in each layer increases in accordance with expression (3), so the length of an advantageous mutant front at colony radius $r_2$ is given by $\frac{r_2}{r_1}(1 + (r_2 - r_1)s_1)$.

- *A 3D cylinder.* Again, this geometry was used in [1]. Wild-type cells form a cylinder with a fixed base of area $S$ and grow as a flat front in one direction perpendicular to the base. Single-hit mutants are created at the rate $Su_1$ per layer. If a neutral mutant is generated at layer $r_1$, it creates a clone of width 1. If however the mutant cells have a selective advantage, the clone will have the shape of a cone, with the front area increasing as $(1 + s_1(r_2 - r_1))^2$.

- A 3D range expansion. Finally, assuming that the colony expands as a 3D sphere, the probability to create a mutant in layer $r$ is $4\pi r^2 u_1$. As in a 2D range expansion, a neutral mutant clone created at radius $r$ will on average grow outwards in such a way that the fraction of mutant cells in each layer is constant. This fraction is equal to $\frac{1}{4\pi r^2}$ for a sphere. If a neutral mutant first appeared at layer $r_1$, the area of its clone's growing front at colony radius $r_2$ is given by $\frac{4\pi r_2^2}{4\pi r_1^2} = \frac{r_2^2}{r_1^2}$. For advantageous mutants, the fraction of mutants in each layer again increases by the same factor as in a 3D cylinder, so the front area is given by $\frac{r_2^2}{r_1^2}(1 + s_1(r_2 - r_1))^2$.

Formula (2) summarizes these ideas for different dimensionalities.

**Remark 1.** The 2D flat front expansion described above corresponds to a colony growth in one direction, but it differs from the "true" 1D system that is included in equation (2) and equations below, including scaling laws. The "true" 1D system corresponds to $W = 1$ and thus the width of a mutant clone, even if it is advantageous, is limited to $W = 1$. For this reason, for the $D = 1$ case, expressions (4), (8), and (11) are all independent on fitness advantage.

**Remark 2.** The concept of reproductive fitness is context specific. For the purposes of this study., the amount of increase, from layer to layer, in the number of mutants undergoing a 2D flat expansion, which is given by expression (3), can be viewed as a definition of the selection coefficient.

Now we return to the derivation of the number of mutants in expanding populations. The size of a colony of advantage $s_1$ that originated at $r_1$ and is measured at $r_2$ is given by

$$n^{s_1}(r_1, r_2) = \int_{r_1}^{r_2} a^{s_1}(r_1, \rho)\, d\rho = \sum_{i=0}^{D-1} \frac{(1 - s_1 r_1)^i (D-1)! s_1^{D-1-i}}{r_1^{D-1} i!(D-1-i)!(2D-1-i)} \left( r_2^{2D-1-i} - r_1^{2D-1-i} \right).$$

In the expression above, we represented $(1 + s(\rho - r_1))^{D-1} = \sum_{i=0}^{D-1} \frac{(1-sr_1)^i s^{D-1-s} \rho^{D-1-i}(D-1)!}{i!(D-i-1)!}$ and then performed the integration in $\rho$.

#### 1.2 One-hit mutants

To obtain the total number of mutants of fitness advantage $s_1$ at size $R$, we have

$$Y_1^{s_1}(R) = \int_0^R u_1 M_0(r) n^{s_1}(r, R)\, dr = 2^{D-1} C_D u_1 (D-1)! \sum_{k=0}^{D-1} \frac{R^{2D-k} s_1^{D-1-k}}{k!(D-k)!(2D-k)}. \quad (4)$$

To arrive at this expression, we used

$$(1 - s_1 r)^i = \sum_{k=0}^{i} \frac{(-1)^{i-k} s_1^{i-k} r^{i-k} i!}{k!(i-k)!}. \quad (5)$$

Then we changed the order of summation: $\sum_{i=0}^{D-1}\sum_{k=0}^{i}\ldots = \sum_{k=0}^{D-1}\sum_{i=k}^{D-1}\ldots$, and performed the summation in $i$, followed by the integration in $r$.

Expression (4) can be written out explicitly as follows:

$$\begin{array}{rc} 1D: & Y_1^{s_1}(R) = \frac{1}{2}R^2u_1, \\ 2D: & Y_1^{s_1}(R) = \pi R^3 u_1\left(\frac{2}{3}+\frac{Rs_1}{4}\right), \\ 3D: & Y_1^{s_1}(R) = \pi R^4 u_1\left(1+\frac{4Rs_1}{5}+\frac{2R^2s_1^2}{9}\right). \end{array}$$

A simpler expression is available for neutral mutants:

$$Y_1^0(R) = \frac{2^{D-1}C_D u_1 R^{D+1}}{D+1}. \tag{6}$$

For an advantageous mutant, the leading term (under the assumption that $Rs_1 \gg 1$) is given by

$$Y_1^{s_1}(r) \sim \frac{2^{D-2}C_D u_1 s_1^{D-1} R^{2D}}{D^2}. \tag{7}$$

#### 1.3 Two-hit mutants

The total surface area of single-hit mutants of fitness advantage $s_1$, measured at size $r$, is given by

$$M_1^{s_1}(r) = \int_0^r u_1 M_0(\rho) a^{s_1}(\rho, r)\, d\rho = \frac{C_D u_1 (2r)^{D-1}}{s_1 D}\left((1+s_1 r)^D - 1\right).$$

The total number of double mutants (with fitness advantages $s_1$ and $s_2$ for the first and second hit) is given by

$$\begin{aligned} Y_2^{s_1,s_2}(R) &= \int_0^R u_2 M_1^{s_1}(r) n^{s_2}(r,R)\, dr \\ &= 2^{D-1}C_D u_1 u_2((D-1)!)^2 \sum_{k=0}^{D-1}\sum_{m=0}^{D-1}\frac{s_1^{D-1-m}s_2^{D-1-k}}{m!k!}\frac{R^{3D-k-m}}{(2D-k-m)!(3D-k-m)} \end{aligned} \tag{8}$$

To obtain this expression, similar to the expression for $Y(s_1)$, we used expansion (5), where instead of $s_1$ we have $s_2$. We further used

$$(1+s_1 r)^D - 1 = \sum_{m=0}^{D-1} s_1^{D-m} r^{D-m}\frac{D!}{m!(D-m)!}.$$

The summation in $i$ and $k$ is rearranged as before, and then integration is performed. Expression (8) can be written out explicitly as follows:

$$\begin{array}{rc} 1D: & Y_2^{s_1,s_2}(R) = \frac{1}{6}R^3u_1u_2, \\ 2D: & Y_2^{s_1,s_2}(R) = \pi R^4 u_1 u_2\left(\frac{1}{4}+\frac{R(s_1+s_2)}{15}+\frac{R^2s_1s_2}{72}\right), \\ 3D: & Y_2^{s_1,s_2}(R) = \pi R^5 u_1 u_2\left(\frac{2}{5}+\frac{2R(s_1+s_2)}{9}+\frac{R^2(s_1+s_2)^2}{21}+\frac{R^3s_1s_2(s_1+s_2)}{60}+\frac{R^5s_1^2s_2^2}{405}\right). \end{array}$$

A simpler expression is available for neutral mutants:

$$Y_2^{0,0}(R) = \frac{2^{D-2} C_D u_1 u_2 R^{D+2}}{D+2}. \tag{9}$$

Further, in the presence of a neutral and an advantageous hit of advantage $s$, the leading term (under the assumption that $Rs \gg 1$) is given by

$$Y_2^{s,0}(r) \sim \frac{2^{D-1} C_D u_1 u_2 s^{D-1} R^{2D+1}}{(D+1)D(2D+1)}. \tag{10}$$

#### 1.4 Three-hit and further mutants

Consider all the double-hit mutants with the fitness advantages $s_1, s_2$ resulting from each of the mutations. The total surface area of this population measured at size $r$ is given by

$$M_2^{s_1,s_2}(r) = \int_0^r u_2 M_1^{s_1}(\rho) a^{s_2}(\rho, r)\, d\rho,$$

and the total number of 3-hit mutants at size $R$ is given by

$$Y_3^{s_1,s_2,s_3}(R) = \int_0^R u_3 M_2^{s_1,s_2}(r) n^{s_3}(r, R)\, dr.$$

In general, the total surface at size $r$ of $(n-1)$-hit mutants is calculated as

$$M_{n-1}^{s_1,\ldots,s_{n-1}}(r) = \int_0^r u_{n-1} M_{n-2}^{s_1,\ldots,s_{n-2}}(\rho) a^{s_{n-1}}(\rho, r)\, d\rho,$$

and the total number of $n$-hit mutants at size $R$ is given by

$$\begin{aligned} Y_n^{s_1,\ldots,s_n}(R) &= \int_0^R u_n M_{n-1}^{s_1,\ldots,s_{n-1}}(r) n^{s_n}(r, R)\, dr \\ &= 2^{D-1} C_D \left(\prod_{i=1}^n u_i\right) R^{(n+1)D} \sum_{i_1=0}^{D-1} \cdots \sum_{i_n=0}^{D-1} \frac{\left(\prod_{k=1}^n \frac{(D-1)! s_k^{D-1-i_k} R^{-i_k}}{i_k!}\right)}{(nD - \sum_{k=1}^n i_k)!((n+1)D - \sum_{k=1}^n i_k)} \end{aligned} \tag{11}$$

Note that this expression does not change if we change the order in which the values $s_1, \ldots, s_n$ and $u_1, \ldots, u_n$ are enumerated. There are several interesting special cases. In particular, if $s_k = 0$ for all $k$, the only nonzero contributions come from $i_k = D-1$, and the expression for the neutral $n$-hit mutant numbers simplifies to

$$Y_n^{0,\ldots,0} = \frac{2^{D-1} C_D}{n!(D+n)} \left(\prod_{i=1}^n u_i\right) R^{D+n}. \tag{12}$$

If one or more mutations are advantageous, there is more than one term in the expression for the number of $n$-hit mutants. The terms have consecutive powers of $R$, with $R^{D+n}$ (the neutral contribution) being the lowest power. To find the highest power, assume that $m$ out of $n$ mutations are advantageous. Without loss of generality, we can have $s_1, \ldots, s_m > 0$ and $s_{m+1}, \ldots, s_n = 0$. The nonzero contributions in the sum in expression (11) must contain $i_k = D-1$ for $m+1 \leq k \leq n$. To maximize the power of $R$, we set $i_k = 0$ for $1 \leq k \leq m$. Simplifying, we obtain the fastest growing (in $R$) term in the expression for the $n$-hit mutants with $m$ advantageous hits:

$$Y_n^{s_1,\ldots,s_m,0,\ldots,0} \approx 2^{D-1} C_D \left( \prod_{i=1}^{n} u_i \right) R^{D+n+m(D-1)} \frac{\left( \prod_{k=1}^{m} s_k^{D-1} \right) ((D-1)!)^m}{(n+m(D-1))!(D+n+m(D-1))}. \quad (13)$$

The number of advantageous $n$-hit mutants starts off growing with the same power $R$ as neutral $n$-hit mutants, and only "accelerates" once the total size increases. For example, if all the advantageous mutant types have the same fitness advantage, $s_i = s$ for $1 \leq i \leq m$, then the $n$-hit mutant population starts off growing as $R^{D+n}$, and only when $R$ reaches approximately $1/s$ it starts behaving as an advantageous $n$-hit mutant population, with the growth law switching to $R^{D+n+m(D-1)}$. If fitness advantage values, $s_1 < \ldots < s_m$ all have a different order of magnitude, then the system undergoes multiple switches, first from $R^{D+n}$ to $R^{D+n+1}$ at size $R \sim 1/s_m$, then to $R^{D+n+2}$ at size $R \sim 1/s_{m-1}$, etc.

Finally, we provide the leading term expression for the number of $n$-hit mutants where only a single mutation results in a fitness advantage ($m = 1$ in expression (13)):

$$Y_n^{s_1,0,\ldots,0} \approx 2^{D-1} C_D \left( \prod_{i=1}^{n} u_i \right) R^{2D+n-1} \frac{s_1^{D-1}(D-1)!}{(D+n-1)!(2D+n-1)}. \quad (14)$$

If $s_1 = 0$, then expression (12) is the leading order approximation for the number of $n$-hit mutants. If $s_1 > 0$ (and assuming $Rs_1 \gg 1$) expression (14) approximates the behavior; for $Rs_1 \ll 1$ mutants are essentially neutral, expression (14) gives an underestimation, and equation (12) should be used instead. This can be summarized as follows,

$$Y_n^{s_1,0,\ldots,0} \approx 2^{D-1} C_D R^{D+n} \left( \prod_{i=1}^{n} u_i \right) \max \left\{ \frac{1}{n!(D+n)}, \frac{(Rs_1)^{D-1}(D-1)!}{(D+n-1)!(2D+n-1)} \right\}. \quad (15)$$

This can be also written as

$$Y_n^{s_1,0,\ldots,0} \approx 2^{D-1} C_D R^{D+n} \left( \prod_{i=1}^{n} u_i \right) \begin{cases} \frac{1}{n!(D+n)}, & Rs_1 < K_0, \\ \frac{(Rs_1)^{D-1}(D-1)!}{(D+n-1)!(2D+n-1)}, & Rs_1 > K_0, \end{cases} \quad (16)$$

where

$$K_0 = \left( \frac{(D+n-1)!(2D+n-1)}{n!(D+n)(D-1)!} \right)^{\frac{1}{D-1}}.$$

#### 1.5 Recursive formulas and closed-form expressions for mutant numbers

To summarize, we have derived the following expressions for the numbers of mutants in expanding populations: formula (4) for single-hit mutants, formula (8) for double-hit mutants, and formula (11) for $n$-hit mutants.

Closed-form expressions are available for neutral mutants (expression (6) for 1-hit mutants, (9) for 2-hit mutants, and (12) for $n$-hit mutants). For advantageous mutants, the fastest (in $R$) growing terms are given by (7), (10), and (13).

While for single-hit advantageos mutants, the sum in (4) is difficult to evaluate, taking an $R$-derivative gives a simpler expression, resulting in the following equation:

$$\frac{dY_1^{s_1}}{dR} = \frac{(2D)^{D-1}u_1}{s_1 D}\left((1+Rs_1)^D - 1\right).$$

This expression is also valid in the limit of small advantage. Taking the limit $s_1 \to 0$, we obtain

$$\frac{dY_1^0}{dR} = 2^{D-1}R^D u_1 = Du_1 Y_0,$$

where we used expression for the wild-type population (1). The above relationship generalizes to an arbitrary number of hits, by taking the $R$ derivative in equation (12):

$$\frac{dY_n^{0,\ldots,0}}{dR} = \frac{D+n-1}{n} u_n Y_{n-1}^{0,\ldots,0}, \quad n = 1, 2, \ldots. \tag{17}$$

This relationship describes the production of multiple-hit neutral mutants in a spatially growing population. Since the colony radius $R$ increases linearly with time, the derivative on the left can be interpreted as a time-derivative. The chain (17) can be constrasted to the usual cascade of mutations formulated for exponentially growing populations:

$$\frac{dy_n}{dt} = u_n y_{n-1} + y_n,$$

where $y_n$ is the population of $n$-hit neutral mutants.

### 2 The role of mutation acceleration: the basic model

Equations (4) and (8) can be used to evaluate the relative contributions of different mutant pathways.

| Parameter | Notation |
|---|---|
| Basic mutation rate | $\mu$ |
| Enhanced mutation rate for accelerated cells | $\mu_A = a\mu \geq \mu$ |
| Rate of acceleration acquisition | $p = b\mu \geq \mu$ |

Table 1: Some notations for the model with mutation acceleration.

Refer to table 1 and figure 1 of the main text for notations. Our theoretical results provide estimates for the different cell subpopulations. In particular, the population of single-hit target mutants, $Z_m$, is given by expression (4) with $s_1 = 0$ and $u_1 = \mu$. The population of double-hit mutant obtained by first acquiring characteristics of an accelerated type and then the target mutation, type $Z_{am}$, is given by expression (8) where $u_1 = p, u_2 =$

$\mu_A, s_1 = s, s_2 = 0$. The population of double-hit mutant where the order of mutations is reversed, $Z_{ma}$, is given by expression (8) where $u_1 = \mu, u_2 = p, s_1 = 0, s_2 = s$.

Our goal is to determine whether the presence of a mutator contributes significantly to the rate of mutant production. To this end, we compare the contribution coming from the double-hit mutants in the presence of acceleration (types $Z_{ma}$ and $Z_{am}$), with the number of mutants obtained in the absence of acceleration ($Z_m$). Denote by $R = R_c$ the solution of the equation

$$Z_m = Z_{am} + Z_{ma}. \tag{18}$$

For $R > R_c$, mutation acceleration makes a significant (100% or more) contribution to the mutant pool.

#### 2.1 Model applicability

The model holds as long as the number of mutants is much smaller than the total number of cells.

**Single mutations.** If follows from equations (6) and (7) that the number of neutral mutants grows as $R^{D+1}$ and the number of advantageous mutants grows as $R^{2D}$, whereas the total number of cells increases as $R^D$ (see equation (1)). Therefore, as the system increases in size, the number of neutral or advantageous mutants will inevitably catch up with the number of wild-type cells, at which point the model is no longer valid. To determine this threshold of model validity, let us denote by $R = R_v$ the solution of the equation $Z_1(R) = Z_{wt}(R)$. For neutral mutants, we use equation (6) and obtain

$$\frac{DR^{D+1}u_1}{D+1} = R^D,$$

resulting in

$$R_v = \frac{D+1}{Du_1}. \tag{19}$$

For advantageous mutations, let us assume that

$$s \gg u_1, \tag{20}$$

and also $u_1 \ll 1$. Then, using expression (7), we obtain

$$R_v = \sqrt{\frac{2(D+2)}{Dsu_1}} \left(1 + O\left(\sqrt{\frac{u_1}{s}}\right)\right). \tag{21}$$

As expected, the model breaks down for smaller values of $R$ if the mutant is advantageous. Plugging in realistic parameters for the mutation rates, we can see that the model holds for a very large range of colony sizes, both for advantageous and neutral mutations.

**Double mutations: the neutral case.** We will use equation (18) to study the model applicability in the case of a system with accelerated types. The expression for $Z_m$ is given by equation (6) with $u_1 = \mu$; the expression for $Z_{am}$ by (9) with $u_1 = b\mu, u_2 = a\mu$; and the expression for $Z_{ma}$ by (9) with $u_1 = \mu, u_2 = b\mu$. Equation (18) gives

$$R_c = \frac{2(D+2)}{(D+1)(a+1)b\mu}. \tag{22}$$

For sizes of the order of $R_c$ or larger, the contribution of mutation acceleration is large. On the other hand, the model is applicable for sizes lower than $R_v$ in equation (19) with $u = b\mu$ (as we are comparing the number of accelerated cells with the total population):

$$R_v = \frac{D+1}{Db\mu}.$$

Will prediction (22) be achieved within the applicability of the model? We have $R_c < R_v$ as long as

$$a > 1 - \frac{2}{(D+1)^2},$$

which definitely holds for $a \geq 1$.

**Double mutations: the avantageus case.** Next, we assume that the accelerated cells are advantageous, such that inequality (20) holds. In this case, $Z_m$ is again given by expression (6) with $u_1 = \mu$, but the expressions for double-mutants are different: $Z_{am}$ is given by expression (10) with $u_1 = b\mu, u_2 = \bar{\mu} = a\mu$, and $Z_{ma}$ by expression (10) with $u_1 = \mu, u_2 = p = b\mu$, $s_1 = 0$ and $s > 0$. Note that these expressins are a good approximation if $Rs \gg 1$, a condition that will be confirmed after a solution is found. Solving equation (18) for $R$, we obtain

$$R_c = \left(\frac{D(2D+1)}{(a+1)b\mu s^{D-1}}\right)^{1/D}. \tag{23}$$

Consequently, $R_c s \gg 1$ because of the assumption (20). As the system approaches sizes of the order of $R_c$ or larger, the contribution of accelerated cells becomes large. In the case of advantageous accelerations, the model is applicable for sizes lower than $R_v$ in equation (21) with $u = b\mu$:

$$R_v = \sqrt{\frac{2(D+2)}{bD\mu s}}.$$

Will prediction (23) be achieved within the applicability of the model? For the case of 2D, we have

$$\left(\frac{R_c}{R_v}\right)^2 = \frac{5}{2(a+1)},$$

such that $R_c < R_v$ as long as $a > \frac{3}{2}$, a condition that holds for all biologically relevant cases.

In the 3D case, we have

$$\left(\frac{R_c}{R_v}\right)^3 = \frac{63}{10(1+a)}\sqrt{\frac{3}{10}}\sqrt{\frac{ub}{s}}.$$

Again, this condition is easily satisfied under inequality (20).

#### 2.2 Fitting numerical data

Spatial stochastic simulations were performed for a number of parameter combinations, for the system depicted in the schematic of figure 1 of the main text, panel (b). Both 2D and 3D systems were considered, and the parameters were varied according to Table 2. In particular, both neutral and advantageous mutations were studied, and parameter $\sigma$ in table 2 denotes the relative increase in the rate of reproduction for all mutants that contained the acceleration mutation. Most systems had zero death rates except for system (b). The basic mutation rate was assumed to be high ($\mu = 10^{-4}$) in order to be able to observe a nontrivial number of double-hit mutants at relatively low population sizes (for computational feasibility). The simulations were run until the population reached the size of about $2 \times 10^7$. The number of independent simulations per parameter combination varied between 3,612 and 51,600.

For each parameter set, the mean trajectory (that is, the average number of cells at different time points) for each of the five populations was found. We denote those numerically obtained values by $Z_{wt}^{num}$, $Z_m^{num}$, $Z_a^{num}$, $Z_{am}^{num}$ and $Z_{ma}^{num}$. Further, the mean total cell number, $V^{num}$, was determined. Then the colony radius was calculated from $V = \pi R^2$ and $V = (4/3)\pi R^3$ in 2D and 3D, respectively. The mean trajectories as functions of the radius are plotted by dots in figure S1.

The data for all four mutant populations were fitted by using functions $KZ_m$, $KZ_a$, $KZ_{am}$, and $KZ_{ma}$, where $K$ is an unknown multiplicative parameter. This parameter (which is of the order of one) serves to create a better match between the scaling laws derived here and the observed mean numbers of cells. It has been noted before (see e.g. [1]) that the scaling laws of this type do not take into account subtle features of the population growth related to the grid and neighborhood structure (e.g. the Moore vs Von Neuman neighborhood), or the effects of cell death. For this reason, we allowed an extra degree of freedom in the fitting function. The second parameter that we fit is the fitness advantage, $s$. The logarithms of the four mutant population sizes were fitted by using our two-parametric functions, and the results are presented by the solid lines in figure S1. The best fit parameters are shown columns $K$ and $s$ of table 2.

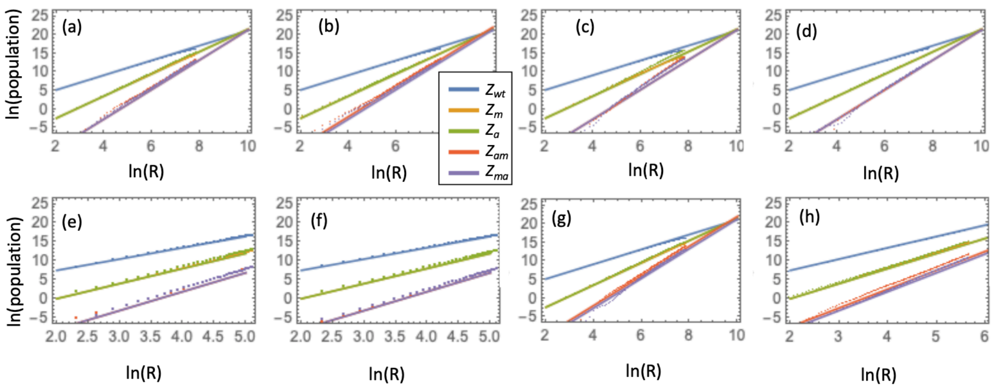

Figure S1: Numerical data (dots) and the best fitting model (lines) of the five populations, as described by the color code on the right of panel (c). The natural logarithms of the cell populations are plotted against the natural logarithms of the colony radius. Simulation parameters are given in table 2.

| System | Dimensionality | $a$ | $d/r$ | $\sigma$ | $K$ | $s$ | # simulations |
|---|---|---|---|---|---|---|---|
| (a) | 2D | 1 | 0 | 0.01 | 1.27 | 0.0008 | 5,160 |
| (b) | 2D | 2 | 0.11 | 0 | 1.43 | 0.0 | 4,644 |
| (c) | 2D | 1 | 0 | 0.03 | 0.94 | 0.005 | 5,160 |
| (d) | 2D | 1 | 0 | 0 | 1.10 | $6 \times 10^{-5}$ | 3,612 |
| (e) | 3D | 1 | 0 | 0.01 | 1.64 | 0.0026 | 51,600 |
| (f) | 3D | 1 | 0 | 0 | 1.52 | $2 \times 10^{-8}$ | 51,600 |
| (g) | 2D | 2 | 0 | 0.01 | 1.10 | 0.0009 | 3,612 |
| (h) | 3D | 2 | 0 | 0.01 | 1.56 | 0.002 | 5,160 |

Table 2: Parameter values used in the simulations of figure S1. In all cases, $\mu = p = 10^{-4}$.

##### 2.3 Nonspatial (exponential) growth

In order to perform a comparison between spatial and non-spatial systems, we modeled mutant growth in a mass-action system by using the usual ordinary differential equations. In particular, for a system without accelerated cells (main text figure 1(a)) it is given by

$$\dot{x}_0 = (1-\mu)x_0, \tag{24}$$

$$\dot{y}_0 = \mu x_0 + y_0, \tag{25}$$

$$x_0(0) = 1, \quad y_0(0) = 0, \tag{26}$$

where $x_0$ and $y_0$ are the wild-type and target mutant populations in the absence of accelerating mutations. For the system with accelerating mutations (main text figure 1(b)) , we have:

$$\dot{x} = (1-\mu)(1-p)x, \tag{27}$$

$$\dot{y}_m = \mu(1-p)x + (1-p)y_m, \tag{28}$$

$$\dot{y}_a = p(1-\mu)x + (1+s)(1-\mu_A)y_a, \tag{29}$$

$$\dot{y}_{ma} = \mu p x + p y_m + (1+s)\mu_A y_a + (1+s)y_{ma}, \tag{30}$$

$$x(0) = 1, \quad y_m(0) = y_a(0) = y_{ma} = 0. \tag{31}$$

Here the populations are denoted by $x$ for wild-type cells, $y_m$ for single-hit mutants with the target mutation, $y_a$ for accelerated cells with no further mutations, and $y_{ma}$ for accelerated cells that contain the target mutation. In systems (24-26) and (27-31), the growth rate of cells is one; this assumption does not make a difference because we are interested in studying mutant numbers as a function of the colony size, and not as a function of time.

Denote by $T_{N,0}$ and $T_N$ the time when the total population reaches size $N$ in the two systems, respectively: $T_{N,0} = \ln N$, $x(T_N) + y_m(T_N) + y_a(T_N) + y_{ma}(T_N) = N$. In figures S3, S4, S5, and similar figures of the main text, the relative contribution of accelerated cells in exponentially growing colonies was calculated as

$$Q = y_{ma}(T_N)/y_0(T_{N,0}). \tag{32}$$

Figure S2 shows that in the nonspatial system, the contribution of duplcated ($\mu_A = 2\mu$) cells is negligible, unless they enjoy a significant advantage. For example, for the parameters of figure S2, a 10% or a 20% advantage leads to only a very small (less than 2%) contribution to the number of mutants.

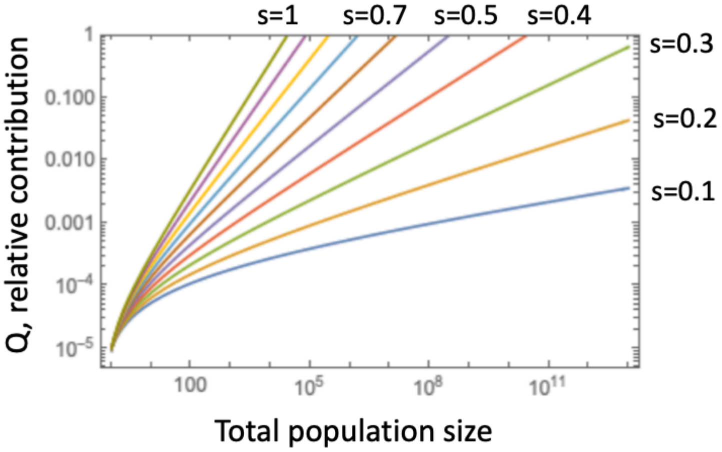

Figure S2: The non-spatial system: the dependence of $Q$ on the advantage of the duplicated cells. basic mutation rate, in the case of gene duplications. The relative contribution of the accelerated type to the total number of mutants, $Q$ (equation (32)), is plotted as a function of the total population size, for the mutant advantage increasing from 10% ro 100%, as indicated by the lines of different colors. The other parameters are $\mu = 10^{-7}$, $\mu_A = 2\mu$, and $p = 100\mu$.

##### 2.4 The relative contribution of accelerated cells: additional results

In addition to the results shown in figure 4 of the main text, here we present further explorations of the parameter space. Figures S3, S4, and S5 explore the parameter dependence of the quantity $Q$, which is the relative contribution of the accelerated type into mutant production. In particular, figure S3 asumes that the acceleration results from gene duplication with $\mu_A = 2\mu$, and varies the basic mutation rate, $\mu$, which is $\mu = 10^{-9}$ in the top row and $\mu = 10^{-5}$ in the bottom row. We can see that in all the cases, the relative contribution of the accelerated type into mutant production is significantly higher for spatial compared to the nonspatial system.

Figures S4 and S5 assume that the mutation acceleration is a hundred-fold, $\mu_A = 100\mu$, and show the results for $Q$ under the assumption that $\mu = 10^{-7}$ and $\mu = 10^{-9}$ respectively.

#### 3 Further models of acceleration

##### 3.1 Two hits to accelerators

Consider the mutation-selection network in figure 5(b) of the main text. We assume that an inactivation of two genes gives rise to an accelerated type (e.g. through the loss of

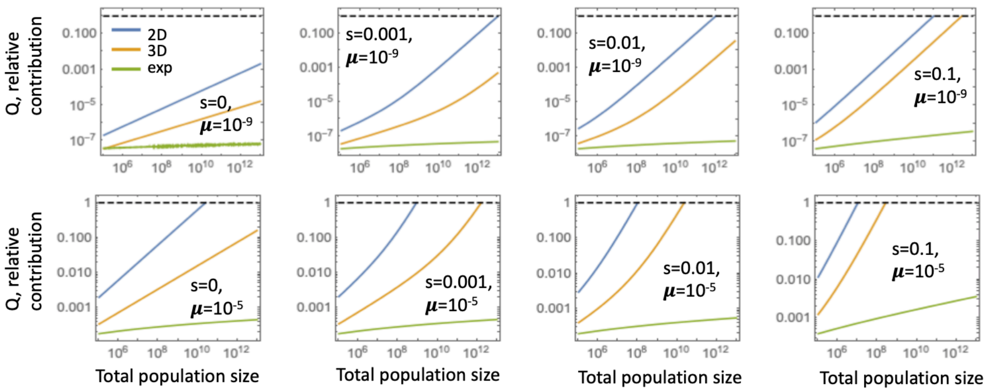

Figure S3: The dependence of $Q$ on the basic mutation rate, in the case of gene duplications. The relative contribution of the accelerated type to the total number of mutants, $Q = (Z_{am} + Z_{ma})/Z_m$ (as predicted by the model), is plotted as a function of the total population size. The three lines in each panel correspond to the 2D (blue), 3D (orange), and mass-action (green) systems. The accelerated mutation rate is $\mu_A = 2\mu$ (duplications). Top row, $\mu = 10^{-9}$. Bottom row, $\mu = 10^{-5}$. The fitness parameter, $s$, increases from left to right as $s = 0, s = 0.001, s = 0.01$, and $s = 0.1$. The dashed horizontal line represents $Q = 1$. In all plots, $p = \mu$.

mismatch repair). The corresponding cell type is denoted as $Z_{12}$, and it is characterized by an accelerated muattion rate $\mu_A$, at which the target mutation can be acquired, creating type $Z_{12m}$. Alternatively, the mutations can be acquired in a different order, giving rise to six alternative pathways from $Z_{wt}$ (the wild type) to $Z_{12m}$ (a mutated, accelerated cell). The contribution of each of these pathways is given by expression (11) with $n = 3, m = 1$; at most one of the values of $s_i$ is positive, which happens if accelerated cells are advantageous. Otherwise, we have $s_1 = s_2 = s_3 = 0$. The only difference among the 6 pathways is the mutation rates of the three steps. Specifically, in 5 out of 6 pathways, two mutation rates are $p$ and the third is $\mu$. In the remaining pathway that includes type $Z_{12}$, the mutation rates are $p$, $p$, and $\mu_A$. Therefore, the analytical approximation for $Z_{12m}$ is given by equation (11), where the product of mutation rates is replaced with $p^2(5\mu + \mu_A)$. The expression for $Z_1$ is given by equation (4) with $s_1 = 0, u_1 = \mu$.

Figure S6 shows the relative contribution of mutation acceleration, $Q = \frac{Z_{12m}}{Z_m}$. In the absence of fitness advantage of the accelerated type ($s_1 = 0$), its contribution is relatively small, especially in 3D. In the presence of fitness advatage, accelerated types make a sigificant conribution to mutation generation for a wide range of parameters. It is intuitively clear that increasing the degree of acceleration, $a = \mu_A/\mu$ and the rate at which accelerated types are generated, $p$, will lead to larger values of $Q$. It is possible to derive simple expressions that characterize how the contribution of acceleration depends on the parameters. To do this, we can solve the equation $Z_m(R) = Z_{12m}(R)$ for $R$, using the leading order terms for $Z_{12m}$ given by equation (15) or (16) with $n = 3$.

The solution of the equation $Z_m(R) = Z_{12m}(R)$, $R = R_c$, gives the colony size when the two contributions become equal. The quantity $R_c$ depends on the combination $\xi = ap^2$. Let us denote the corresponding colony volume (population size) by $V_c$. We have the following result for the colony size, at which acceleration makes a contribution to mutant generation

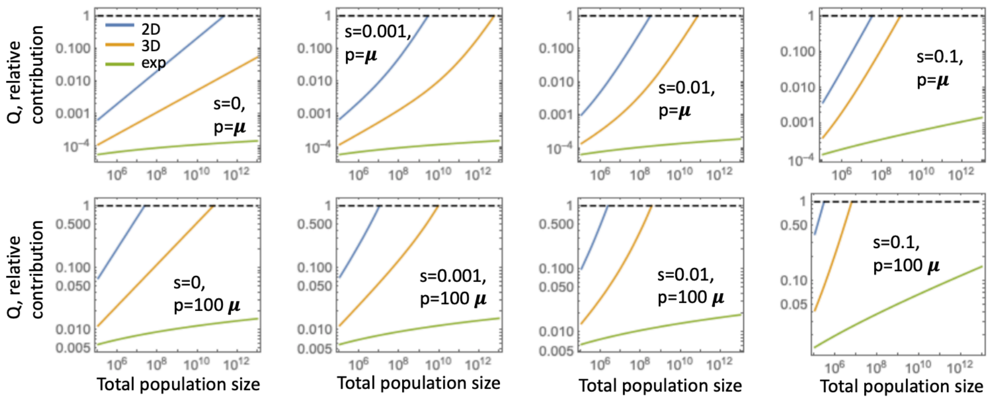

Figure S4: An alternative acceleration assumption: $\mu_A = 100\mu$. The relative contribution of the accelerated type to the total number of mutants, $Q = (Z_{am} + Z_{ma})/Z_m$ (as predicted by the model), is plotted as a function of the total population size. The three lines in each panel correspond to the 2D (blue), 3D (orange), and mass-action (green) systems. Top row, $p = \mu$; bottom row, $p = 100\mu$. The fitness parameter, $s = 0, s = 0.001, s = 0.01$, and $s = 0.1$. The dashed horizontal line represents $Q = 1$. In all plots, $\mu = 10^{-7}$.

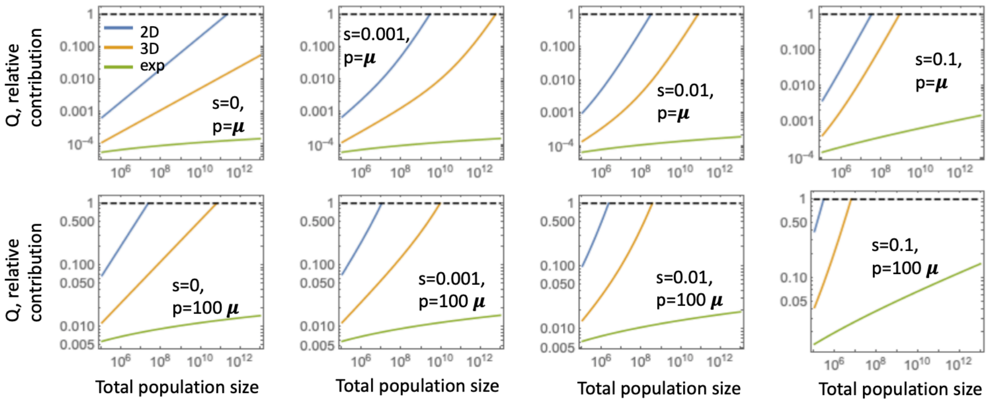

Figure S5: Same as figure S4, except in all plots, $\mu = 10^{-9}$.

that is comparable to direct mutations:

$$2D: \quad V_c = \min\left\{C_{0,2}\xi^{-1}, C_{s,2}(\xi s_1)^{-2/3}\right\}, \tag{33}$$

$$3D: \quad V_c = \min\left\{C_{0,3}\xi^{-3/2}, C_{s,3}(\xi s_1^2)^{-3/4}\right\}, \tag{34}$$

where $\xi = ap^2$ and the constants are given by

$$C_{0,2} = 10\pi \approx 31.4, C_{0,3} = 36\pi \approx 113.1, C_{s,2} = 4\times 6^{2/3}\pi \approx 41.5, C_{s,3} = 16\times(2/3)^{1/4}5^{3/4}\pi \approx 151.2.$$

#### 3.2 A cascade of amplifications

The second mutation-selection network that we consider here is given in figure 5(a) of the main text. In this system we assume that at a rate $p$, a gene duplication event may happen,

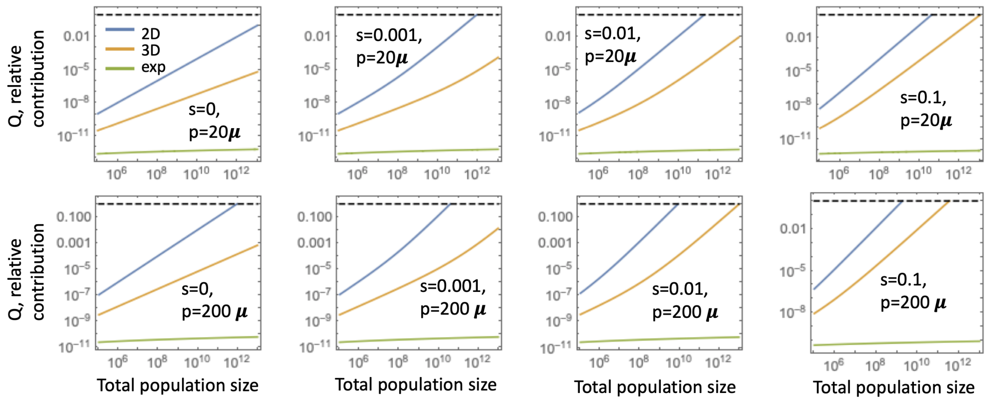

Figure S6: Two hits to acceleration (see selection-mutation diagram of figure 5(b) of the main text). The relative contribution of the accelerated type to the total number of mutants, $Q = Z_{12m}/Z_m$ (as predicted by the model), is plotted as a function of the total population size. The three lines in each panel correspond to the 2D (blue), 3D (orange), and mass-action (green) systems. Top row, $p = 20\mu$; bottom row, $p = 200\mu$. The fitness parameter, $s = 0, s = 0.001, s = 0.01$, and $s = 0.1$. The dashed horizontal line represents $Q = 1$. In all plots, $\mu = 10^{-7}$ and $\mu_A = 100\mu$.

which gives rise to the type denoted by $Z_1$. Further amplification events happen (at rate $p$) giving rise to types $Z_2$, $Z_3$, etc. All the types $Z_k$ with $k = 1, 2, \ldots$ are accelerated to an increasing degree, and each is characterized with an enhanced mutation rate, $\mu_{Ak}$, with $\mu_{Ak} > \mu_{Aq}$ as long as $k > q$. The types $Z_{1m}, Z_{2m}, \ldots$ are produced through mutation acceleration, and the relative contribution of accelerated types is given by

$$Q = \frac{\sum_{k=1}^{\infty} Z_{km}}{Z_m}.$$

Each of the accelerated types containing the target mutation can be created by means of multiple pathways. For type $Z_{km}$ with $k = 1, 2, \ldots$, there are $k+1$ mutation events that separate them from $Z_{wt}$. These pathways to $Z_{km}$ contain a single "vertical" step (corresponding to a target mutation) and $k$ "horizontal" steps with rate $p$, which are amplification events. There are $k+1$ such pathways, and the products of the mutation rates along these pathways are

$$p^k\mu, \; p^k\mu_{A1}, \; \ldots, p^k\mu_{Ak}.$$

The abundance of cells of each accelerated type containing the target mutation, $Z_{km}$, is given by equation (11) with $n = k+1$ and with the product of mutation rates replaced by the sum of the mutation rates over the $k+1$ pathways,

$$\mu_k^{tot} = p^k \left( \mu + \sum_{i=1}^{k} \mu_{Ai} \right) = p^k \mu B_k,$$

where $B_k = 1 + \frac{1}{\mu} \sum_{i=1}^{k} \mu_{Ai}$ depends on the modeling assumptions about the increase in the mutation rates in the amplification cascade.

While there are many possibilities, below we will consider two examples:

- A linear increase model: $\mu_{Ak} = (k+1)\mu$, that is, a single amplification event results in a doubling of the mutation rate, the next one triples it, etc. In this case, $B_k = \frac{(k+1)(k+2)}{2}$.
- A geometric increase model: $\mu_{Ak} = 2^k\mu$, that is, a single amplification event results in a doubling of the mutation rate, the next one quadruples it, etc. In this case, $B_k = \left(2^{k+1} - 1\right)$.

**The neutral case.** Let us first consider the system where the amplification events are neutral. The quantity $Z_{km}$ is given by equation (12) with $n = k + 1$, where the product of mutation rates is replaced by $\mu_k^{tot}$. We have

$$\sum_{k=1}^{\infty} Z_{km} = 2^{D-1} C_D R^{D+1} \mu \sum_{k=1}^{\infty} (Rp)^k \frac{B_k}{(k+1)!(D+k+1)}.$$

While a closed form expression is available for this sum, it is enough to note that in biological systems considred here, the quantity $Rp \ll 1$, and thus the contribution of the amplification cascade beyond the first duplication event is negligible.

**Amplified cells have the same, enhanced, fitness.** Now, equation (14) gives the leading order term in the growth of the mutant (again, we take $n = k + 1$ and replace the product of mutation rates by $\mu_k^{tot}$). We have

$$\sum_{k=1}^{\infty} Z_{km} = 2^{D-1} C_D (D-1)! R^{2D} \mu s_1^{D-1} \sum_{k=1}^{\infty} (Rp)^k \frac{B_k}{(D+k)!(2D+k)}.$$

As in the neutral case, $Rp \ll 1$, and the contribution of the amplification cascade beyond the first duplication event is negligible.

**Each consecutive amplification increases fitness.** Finally, we consider the case where each amplificaion event results in a fitness gain compared to the previous type. For simplicity, here we assume that the gain is given by the same value, $s$, after each amplification (more general situations can be considered in a similar fashion). In this case, we will use expression (13) with $n = k + 1, m = k$ and the product of mutation rates replaced by $\mu_k^{tot}$; this gives the highest order term in each contribution. We have,

$$\sum_{k=1}^{\infty} Z_{km} = 2^{D-1} C_D R^{D+1} \mu \sum_{k=1}^{\infty} (R^D s^{D-1} p)^k \frac{B_k}{(kD+1)!((k+1)D+1)}.$$

Now, the consecutive terms on the right hand side multiply powers of $(R^D s^{D-1} p) \sim Z_{wt} p s^{D-1}$. This parameter combination can be non-small in realistic scenarios.

Figure S7 shows a comparison between target mutations generated directly ($Z_m$, black dashed lines), target mutations generated by duplicated cells ($Z_{1m}$, red dashed lines), and target mutations generated by all the amplified cells in the cascade ($\sum_{k=1}^{\infty} Z_{km}$, red solid lines). Mutant numbers are shown as functions of the total population size in 2D (a) and 3D (b), for different values of the fitness advantage, $s$. It is assumed that each consecutive

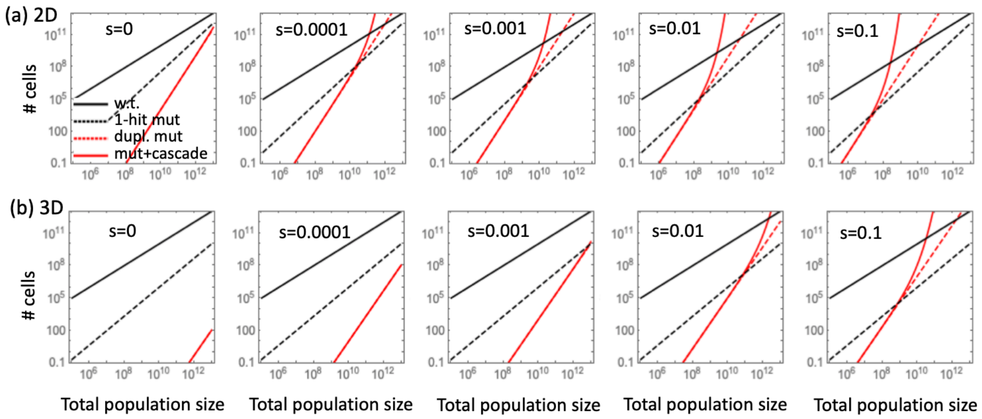

Figure S7: A cascade of amplifications. The number of mutants produced by amplified cells is shown as a function of the colony size, for a (a) 2D and (b) 3D system. Red solid lines correspond to the number of target mutations obtained through the cascade of duplications ($\sum_{k=1}^{\infty} Z_{km}$), while red dashed lines contain only duplicated mutant cells ($Z_{1m}$). For comparison, black dashed lines are single-hit target mutants ($Z_m$), and black solid lines are wild type cells. The selection coefficient varies (and affects equally each conscutive amplification event). The rest of the parameters are $\mu = 10^{-7}, p = 100\mu$; a geometric copy increse model is used.

amplification increases fitness, and that the copy number increases geometrically. The number of wild-type cells is shown by solid black lines, and it should be noted that the model breaks down when any mutant numbers approach the number of wild type cells.

We observe that, as predicted, the existence of further amplification events in the cascade increases the number of mutants generated in the system, compared to the number of mutants generated by duplicated cells only (given that each duplication event confers an incremental fitness advantage). The effect is seen for larger colony sizes and larger values of $s$, and is more pronounced in 2D compared to 3D.

Note that the summation over an infinite number if types is obviously a mathematical abstraction. In a realistic system, only a finite number of types (say, $n$ types) are characterized by nonnegative values of $s$, while all subsequent types are disadvantageous. In this case, the relevant quantity is $\sum_{k=1}^{n} Z_{km}$.

### 4 An example of a model of fitness in a 2D flat expansion

First assume the Wright-Fisher model, with the constant population of size $N$ and non-overlapping generations. Suppose that currently, there are $m$ mutants and $N - m$ wild type cells. In the next generation, the population is created by sampling with replacement, such that for each sampling, the probability to pick a mutant is given by

$$\frac{m(1+s)}{m(1+s)+N-m} \approx \frac{m(1+s)}{N},$$

assuming $s \ll 1$. After $j$ generations, the mutant fraction increases by a factor of $(1+s)^j \approx 1+js$. In our application, the generations of the Wright Fisher process play the role of the consecutive layers of the expanding colony, which, starting from a single mutant, are expected to have

$$1, 1+s, 1+2s, 1+3s, \ldots, 1+js$$

mutants.

This process is non-spatial. To take account of the spatial effects, consider the example of a 2D flat front expansion. Suppose that $m > 2$ mutants occupy adjacent spots, and the rest of the individuals are wild-type. When the next layer is created, we assume that the individual in each position is sampled from its three neighbors (in the Moore neighborhood) that belong to the previous layer, see figure S8. Exactly $m-2$ spots in the next layer (such as cell "C") have only mutant neighbors in the existing layer, and will therefore be occupied with mutants. Spots that only have wild type neighbors (such as cell "D") will be occupied by wild types. The only positions that have a possibility to be occupied by either mutants or wild types are spots marked with "A" and "B", that is, the four spots that are adjacent to both mutant and wild type cells at the interface. For the two cells denoted by "A", the competition is between two wild type cells and one mutant cell. The probability that spot "A" is occupied by a mutant is given by

$$P_A = \frac{1+s}{1+s+2} \approx \frac{1}{3} + \frac{2s}{9}.$$

For the two cells denoted by "B", the competition is between two mutant cells and one wild type cell. The probability that spot "B" is occupied by a mutant is given by

$$P_B = \frac{2(1+s)}{2(1+s)+1} \approx \frac{2}{3} + \frac{2s}{9}.$$

It is possible that out of the 4 spots (denoted by "A" and "B"), $j$ spots will contain a

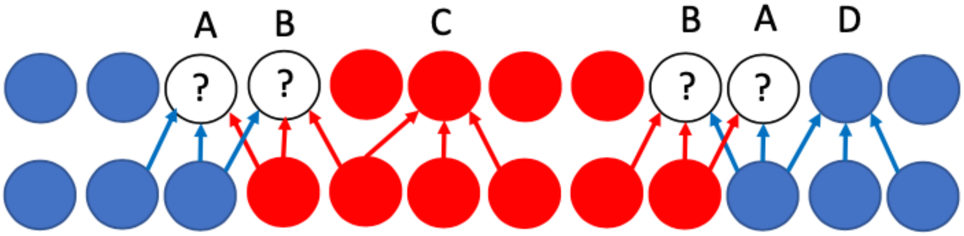

Figure S8: A schematic showing the process of layer creation in the case of 1D flat expansion. Mutant cells are shown by red circles and wild type cells by blue circles. The left column is the existing layer and the right column is the next layer. Arrows pointing toward a cell indicate which cells compete to occupy that spot. The question mark denotes spots where the type is uncertain. Different position types are denoted by "A", "B", "C", and "D", and described in the text.

mutant, with $0 \leq j \leq 4$. We can calculate the probability $P(j)$ that exactly $j$ cells are mutant:

$$P(0) = (1-P_A)^2(1-P_B)^2, \quad P(1) = 2P_A(1-P_A)(1-P_B)^2 + 2(1-P_A)^2P_B(1-P_B),$$
$$P(2) = P_A^2(1-P_B)^2 + P_B^2(1-P_A)^2 + 4P_AP_B(1-P_A)(1-P_B),$$
$$P(3) = 2P_AP_B^2(1-P_A) + 2P_BP_A^2(1-P_B), \quad P(4) = P_A^2P_B^2.$$

The expected number of mutants is given by

$$\sum_{j=0}^{4} jP(j) = 2(P_A+P_B) = 2 + \frac{8}{9}s.$$

This means that on average, in each generation, the number of mutants will increase by a constant value $(8s)/9$. Starting from a single mutant, the mean number of mutants in subsequent layers are therefore

$$1, 1+\frac{8}{9}s, 1+2\frac{8}{9}s, 1+3\frac{8}{9}s, \ldots, 1+j\frac{8}{9}s.$$

(Note that this rule is modified when the number of mutants is $m < 2$, but this is not incorporated in the sequence above).

This calculation shows that a spatial structure weakens selection for advantageous mutants. This is because most of them do not have a chance to compete with wild types and behave as if they are neutral, and the gain due to selection only happens at the edges.

### References

[1] Dominik Wodarz and Natalia L Komarova. Mutant evolution in spatially structured and fragmented expanding populations. *Genetics*, 216(1):191–203, 2020.
